# Epidermal Langerhans Cells Drive Painful Diabetic Neuropathy in a Sex-dependent Manner

**DOI:** 10.1101/2025.01.30.635734

**Authors:** Paola Pacifico, Dale George, Nirupa D. Jayaraj, Dongjun Ren, James S. Coy-Dibley, Abdelhak A. Belmadani, Mirna Andelic, Daniele Cartelli, Grazia Devigili, Raffaella Lombardi, Giuseppe Lauria, Amy Paller, Richard J. Miller, Daniela M. Menichella

## Abstract

The interaction between non-neuronal cells and nerve endings in the epidermis significantly influences the development of various diseases, including Painful Diabetic Neuropathy (PDN). PDN is a common and challenging complication of diabetes, characterized by changes in skin innervation accompanied by neuropathic pain. While there is growing evidence that epidermal non-neuronal cells, such as resident immune cells, play a crucial role in the progression of PDN, the underlying mechanisms of this neuropathy remain poorly understood. In our studies, we utilized transgenic methods, pain behavioral assessments, and single-cell RNA sequencing (scRNAseq) in a clinically relevant high-fat diet (HFD) mouse model of PDN and skin biopsy samples from PDN patients, to investigate the role of epidermal Langerhans cells (LCs) in this condition. We observed an increased density of LCs in the skin of PDN male mice coinciding with the onset of mechanical allodynia. Furthermore, we found that LCs density correlated with small fiber degeneration in HFD mice and skin biopsies taken from well-characterized PDN patients compared to healthy controls. Importantly, the selective ablation of LCs using a diphtheria-toxin strategy was found to prevent nociceptive behavior in male mice. This indicates that LCs are both necessary and sufficient for the development of mechanical allodynia and spontaneous pain in HFD male mice. Interestingly, when LCs were ablated in HFD female mice, it did not prevent but rather promoted pain behavior, suggesting the existence of sex-specific mechanisms mediated by LCs. scRNAseq transcriptomic analysis of the paw epidermis from HFD mice, which included both males and females, revealed significant sex-mediated differences in the expression of specific target genes within this PDN model. In male mice, scRNAseq identified differentially expressed genes associated with axonal guidance and immune responses in LCs. Additionally, by integrating single-cell RNA data from the epidermis and dorsal root ganglia (DRG) of male mice, we uncovered altered communication between LCs and cutaneous afferents through Semaphorin-Plexin signaling pathways in PDN. These findings highlight LCs as key contributors to the development of PDN and suggest their potential as therapeutic targets for innovative treatments, particularly topical therapies aimed at modulating immune cell activity and neuroimmune communication in the skin. Our investigations of male and female mice indicate that LCs may play different roles in mechanical sensation under both normal and pathological conditions, such as PDN. This underscores the importance of considering sex differences when developing more effective treatments.

**BRIEF SUMMARY:** The interaction between non-neuronal cells and nerve endings in the epidermis plays a significant role in the development of Painful Diabetic Neuropathy (PDN), a challenging complication of diabetes. Using transgenic methods, pain behavioral assessments, and single-cell transcriptomics in a clinically relevant high-fat diet (HFD) mouse model of PDN, and in skin tissues from PDN patients, we identified epidermal Langerhans cells (LCs) as key contributors to the development of PDN. In these studies, we revealed, for the first time, a dimorphic role for LCs in the development of mechanical allodynia and spontaneous pain in a HFD model of PDN. Single-cell RNA sequencing (scRNAseq) of the paw epidermis from both male and female HFD mice identified sex-mediated differences in the expression of specific target genes within this PDN model. In male mice, scRNAseq of LCs revealed differentially expressed genes related to immune responses and axonal guidance, including Semaphorin-Plexin signaling pathways.

## INTRODUCTION

Painful diabetic neuropathy (PDN) is characterized by neuropathic pain that results from the hyperexcitability of nociceptive neurons in the dorsal root ganglion (DRG)(1, 2). PDN is associated with both the degeneration and regeneration of DRG neuron axons that innervate the skin. This picture constitutes small fiber neuropathy, which is the earliest phenotype of PDN(3–5). One critical barrier to developing effective treatments for PDN is the lack of understanding of the molecular mechanisms that lead to neuropathic pain and to small fiber neuropathy. In healthy subjects, DRG neurons extend their axons into the periphery with cutaneous nerves terminating in the epidermis(6, 7), the outermost stratified layer of the skin, However, in patients with PDN, there is a remodeling of the cutaneous innervation leading to both degeneration and regeneration of cutaneous nerves(3, 8, 9).

The axons of different neuronal subpopulations terminate in the epidermis and make both gap junctions and synapse-like contacts with epidermal non-neuronal cells(10–13). The epidermis is primarily composed of keratinocytes, but it also contains immune cells, such as epidermal Langerhans cells (LCs) and a subtype of T lymphocytes, which act as the first line of immune defense against damage(14–17). Recent evidence underscores the significant role of non-neuronal cells in the skin in the development of PDN(18), with inflammatory processes likely playing a crucial role(19). Inflammatory markers, including interleukins IL-6, IL-2, and tumor necrosis factor-*α* (TNF*α*), are elevated in individuals with diabetes, suggesting a chronic, low-grade inflammatory state in diabetic patients(20) (21, 22). Additionally, macrophage density is increased in the dermis in patients with painful versus non-painful diabetic neuropathy(23).

LCs, first described by Paul Langerhans in 1868(24), are key components of the innate immune response(25). LCs exhibit characteristics of resident antigen-presenting cells, which are responsible for tissue surveillance through the extension and retraction of their dendritic projections(25, 26). They are also ontogenetically related to macrophages(27, 28). Although LCs have been associated with various inflammatory skin diseases, their specific role in PDN remains largely unexplored. As immune cells, LCs not only conduct tissue surveillance but also release and respond to multiple inflammatory molecules(25, 27, 29, 30). This activity may contribute to the elevated levels of cytokines observed in patients with neuropathic pain conditions(18, 20, 31–33). When activated, LCs secrete pro-inflammatory cytokines which could sensitize nociceptor terminals(34, 35). Interestingly, studies have found that LCs density is increased in patients with small fiber neuropathy(34), as well as in a genetic rodent model of type 2 diabetes(36), indicating the potential relevance of these cells in PDN. However, the specific role of LCs in mediating PDN remains unclear.

There is extensive crosstalk between cutaneous afferents and non-neuronal cells in the epidermis, including between keratinocytes and LCs(7). Although mechanisms underlying the involvement of LCs in PDN remain poorly investigated, studies show their interaction with sensory terminal afferents(35, 37), such as Mrgprd-positive nerve terminals(37). From previous work we know that Mrgprd-positive afferents control the recruitment of LCs in the skin(37). Notably, the survival of Mrgprd-expressing nerve fibers depends on their association with LCs(37), suggesting that this neuro-immune communication may be important in mediating neuronal sensitization, as well as axonal degeneration/regeneration in PDN. However, mechanisms promoting the interaction between LCs and Mrgprd-positive afferents remain poorly understood.

In these studies, we demonstrated, for the first time, a dimorphic role for LCs in the development of mechanical allodynia and spontaneous pain in a HFD mouse model of PDN. Single-cell RNA transcriptomic analysis of the paw epidermis from both male and female HFD mice identified sex-mediated differences in the expression of specific target genes within this PDN model. In male mice, single-cell RNA sequencing (scRNAseq) of LCs revealed differentially expressed genes related to axonal guidance and immune responses, including those involved in Semaphorin-Plexin signaling pathways.

## RESULTS

### Epidermal LCs density is increased in the HFD mouse model of PDN at the onset of mechanical allodynia

Accumulating evidence suggests that non-neuronal cells in the skin play a crucial role in the development of PDN(18). While LCs have been implicated in several inflammatory skin diseases, their specific role in PDN has not been extensively explored. Interestingly, increased LCs density has been observed in patients with small fiber neuropathy(34) as well as in the genetic rodent model for type 2 diabetes(36) suggesting that LCs may be relevant in the context of PDN. To investigate this further, we analyzed LCs density in the clinically relevant and well-established HFD model of PDN(38–41). In agreement with the literature(42), we found that feeding a HFD to mice for ten weeks induced obesity, glucose intolerance, pain behaviors such as mechanical allodynia, and remodeling of cutaneous innervation(38, 40–43). In the HFD mouse model of PDN, we observed a dramatic increase of the LCs density in the whole-mount epidermis of wild-type male mice fed HFD for 10 weeks, compared to regular diet (RD) mice **(Fig. 1A-B)**. To determine whether the increase in LCs in HFD mice was localized to the mouse paw epidermis or indicative of more systemic inflammation resulting from the activation and proliferation of LCs, we labeled LCs in whole-mount ears of RD and HFD mice. Our findings showed no differences in LCs density in the ears. **(Suppl. Fig. 1A)**. Thus, similar to PDN patients, where the distal lower extremities are primarily affected, the HFD results in a localized increase in LCs within the epidermis of the mouse paw.

**Figure 1.**
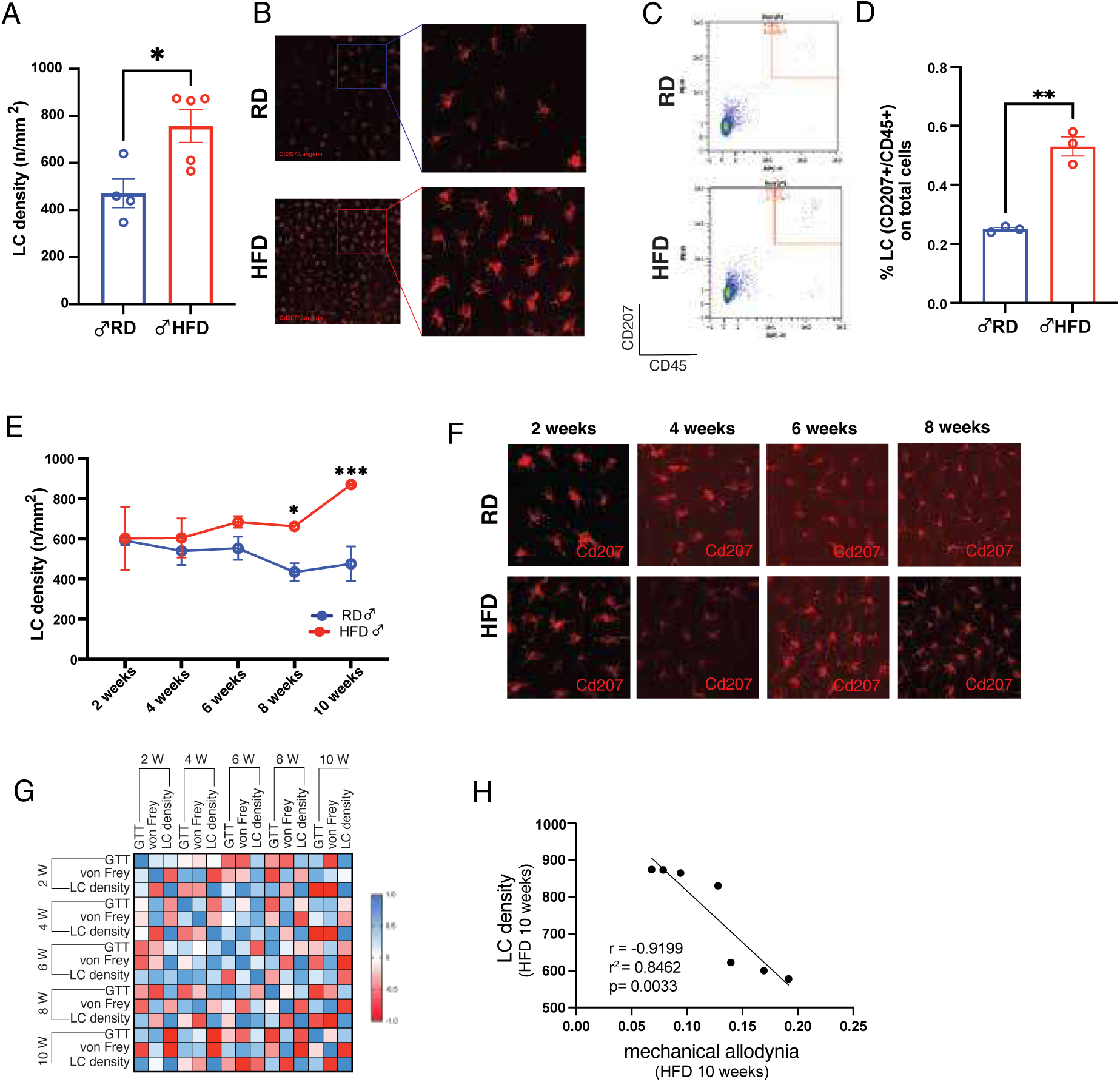
Increased LCs density in HFD male mice. **(A-B)** Quantification of LCs’ density (number of LCs / mm^2^) in RD and HFD male mice at 10 weeks (a) and representative images (b). n=4 animals for RD and n=5 animals for HFD. n=3 sections per animal were acquired and Cd207+ cells per 0.04 mm^2^ per section area were counted. Unpaired two-tailed t-Test with Welch’s correction p=0.0176 (*). **(C)** MACSQuant Tyto cell sorting of Cd207+/Cd45+ gated cells from single cells paw epidermis suspension. PE-Cd207 and APC-Cd45 antibodies were used co-label LCs. **(D)** Graph shows the average percentage from three replicates of Cd207+/Cd45+ sorted cells in RD and HFD. Unpaired two-tailed t-Test p=0.0010 (**). n= 4 animals per diet condition per each replicate. **(E)** Time-course experiment of LCs density at 2, 4, 6, 8 and 10 weeks show the progressive increase of LC in HFD. Two-way ANOVA for multiple comparisons. Between RD and HFD: 8 weeks p=0.0216 (*). 10 weeks p=0.0014 (**). Within HFD: 2w vs 10w p=0.0111; 4w vs 10w p=0.0116; 6w vs 10w p=0.0486; 8w vs 10w p=0.0469. 2weeks: n=3 animals for both RD and HFD. 4 weeks: n=3 animals for both RD and HFD. 6 weeks: n=3 animals for RD and n=4 for HFD. 8 weeks: n=4 animals for RD and n=3 for HFD. 10 weeks: n=4 animals for RD and n=3 for HFD. **(F)** Representative images of LCs Cd207+ in epidermal sheets of RD and HFD mice at different time points. **(G)** Correlation matrix of HFD male features at different time points (2, 4, 6, 8, 10 weeks) including GTT, mechanical allodynia (vonFrey) and LCs density. Red squares indicate negative correlations. 10 weeks vonFrey-LCs: Pearson r = -0.9199, p=0.003. **(H)** Negative correlation between LCs density and mechanical allodynia measured with vonFrey test in 10 weeks HFD mice. Pearson r coefficient -0.9199; r^2^= 0.8462; p value =0.0033.

Next, we wanted to validate these findings using an additional complementary methodology. We elected to use flow cytometry, commonly adopted for isolating and quantifying immune cells, including LCs(37, 44–47). Similarly to histological analysis, flow cytometry showed an increase in the Cd45+/Cd207+ gated cell population confirming the increased number of LCs in HFD male mice **(Fig. 1C-D)**. Given the delicate nature of LCs and the reduced abundance, we employed the gentle MACSQuant Tyto sorter (Miltenyi Biotec, Bergisch-Gladbach, Germany) for cell sorting from single epidermal cell suspensions. We combined three mice for each diet condition and each replicate, illustrating that the percentage of Cd45+/Cd207+ cells in HFD was double that of Cd45+/Cd207+ cells in RD **(Fig. 1D)**.

To investigate the potential contribution of LCs to the development of mechanical allodynia in the HFD mouse model, we measured the LCs density at different time points after starting either RD or HFD (2 weeks, 4 weeks, 6 weeks, 8 weeks and 10 weeks). We observed a gradual increase of LCs over time that became significant starting at 8 weeks, when mice on HFD display mechanical allodynia and small fiber degeneration(40) **(Fig. 1E-F).** We also investigated the relationship between LCs and various features of the HFD phenotype at different time points **(Fig. 1G)**. After ten weeks on an HFD, we found a correlation between the number of LCs and the mechanical threshold in male mice. This suggests that a higher number of LCs is associated with a lower mechanical threshold, indicating the presence of mechanical allodynia **(Fig. 1H).** Similarly, in male mice on a RD, we observed a correlation between the number of LCs and the mechanical threshold, although this was within the range of normal mechanical thresholds in RD mice **(Suppl. Fig. 1B-C).**

### Small fiber neuropathy is associated with LCs density in the epidermis in the HFD mouse model of PDN and in patients with PDN

PDN is characterized by neuropathic pain and the loss or retraction of DRG neuron axons that innervate the skin, also known as small fiber neuropathy(4, 5). In the HFD model of PDN, we have shown that beginning at 8 weeks, mice fed a HFD exhibited a significant reduction in intraepidermal nerve fiber (IENF) density, expressed as the number of nerves crossing the epidermal-dermal junction as a function of length(40). To determine whether LCs density contributes to axonal degeneration in the HFD model, we measured the ratio between the IENFD and LCs density. Our analysis revealed that this ratio was significantly reduced in mice that had been fed an HFD for 10 weeks **(Fig. 2A-B)**. These findings suggest that the loss of cutaneous innervation in the epidermis, a hallmark of small fiber neuropathy, is associated with an increase in LCs density in the epidermis of the HFD mouse model for PDN.

**Fig. 2.**
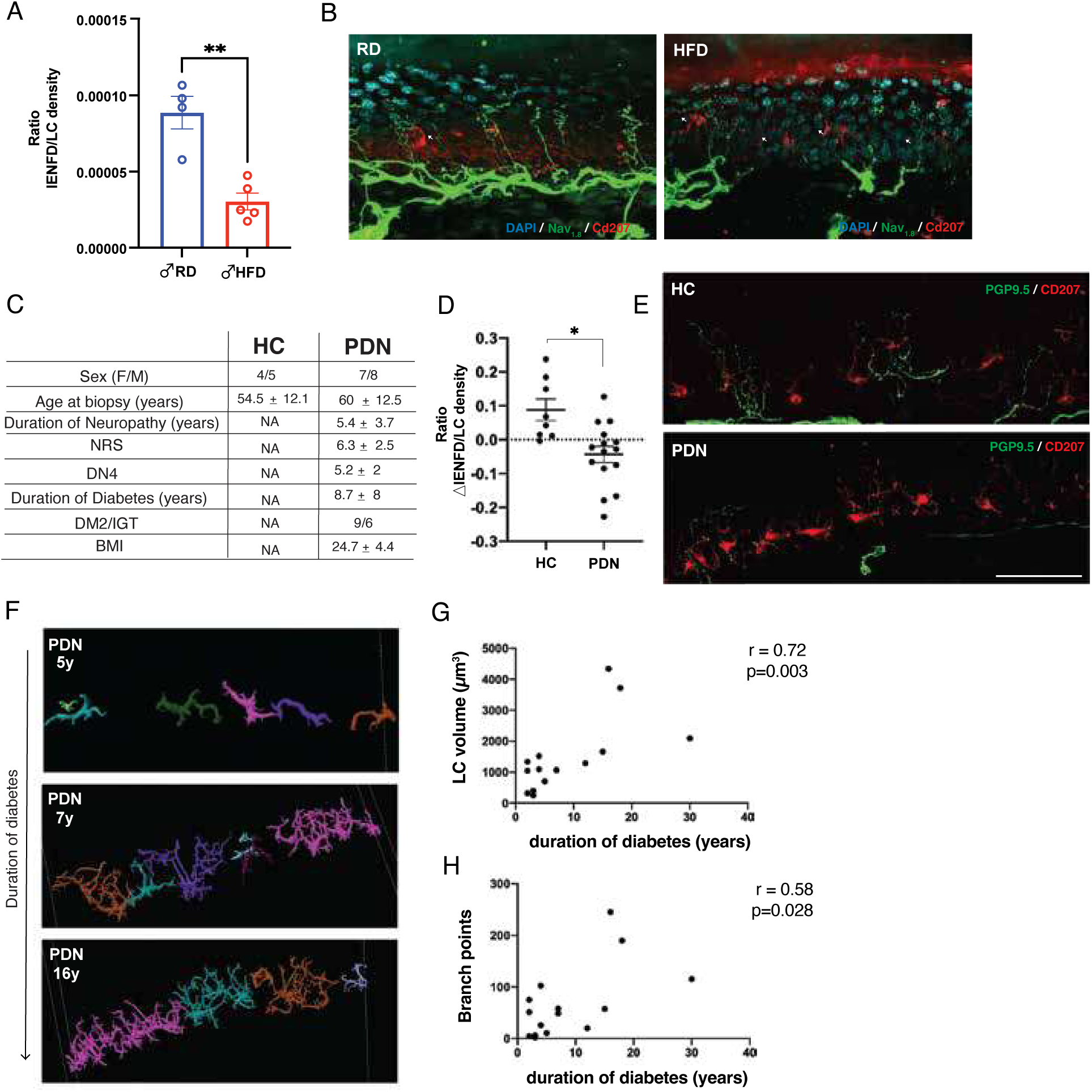
Both HFD mice and PDN patients show reduced cutaneous fibers-LC ratio. **(A)** ratio between intraepidermal nerve fibers density and LCs density is significantly decreased in HFD male mice. Unpaired two-tailed t-Test with Welch’s correction p=0.0066 (**). **(B)** representative confocal images of nociceptive fibers Nav1.8+positive (green) and LCs Cd207+ (red) in skin sections of RD and HFD male mice. **(C)** clinical summary of patients (PDN) and relative healthy controls (HC) involved in the study. **(D)** ratio between the denervation (ΔIENFD), calculated as deviation of the subjects IENFD from the normative value corrected for age and sex, and LCs density in healthy subjects or PDN patients. Mann-Whitney U test. p<0.05 (*). Middle lines of the plots represent the mean value, while whiskers are SEM. **(E)** representative confocal micrographs of epidermal nerve fibers PGP 9.5+ (green) and LCs CD207+ (red) in skin sections of healthy subjects and PDN patients. Scale bar= 50 mm. **(F)** 3D reconstruction of LCs of patients suffering for diabetes for 5 (5 yrs), 7 (7yrs) and 16 (16 yrs) years. **(G)** Scatter correlation plots between LCs volume and duration of diabetes. Spearmann correlation r=0.72, p=0.003 **(H)** Scatter correlation plots of number of branch points and duration of diabetes. Spearmann correlation r=0.58, p=0.028.

To enhance the translational relevance of our studies, we analyzed key parameters from skin biopsies obtained from well-clinically characterized PDN patients and healthy controls. We included 15 PDN patients and 9 healthy controls **(Fig. 2C).** Consistent with previous studies(3, 4, 31, 34, 48–50), we observed a marked reduction of the IENFD in PDN patients compared to controls **(Suppl. Fig. 2A)**. We performed an analysis on these skin biopsies similar to that conducted on HFD mice. We found that the ratio between the fiber denervation (ΔIENFD)—calculated as the deviation of each subject’s IENFD from the normative value adjusted for age and sex; the LCs density was significantly reduced in PDN patients compared to controls **(Fig. 2D-E)**. These findings suggest that the loss of cutaneous innervation in the epidermis is associated with LCs density in the epidermis in the HFD mouse model of PDN and patients with PDN.

Despite the unaltered LCs density in PDN skin sections **(Suppl. Fig. 2B)**, we discovered that both the volume of LCs and the complexity of their arborization processes positively correlated with the duration of diabetes in PDN patients **(Fig. 2F-H).** Specifically, our findings indicated that as the duration of diabetes increased, the size of LCs and the complexity of their branching structures became more pronounced. As in other immune cells which exhibit remarkable plasticity, LCs morphology is linked their function(51), highlighting that potential structural and functional adaptation or compensatory mechanisms within the epidermis occur in PDN. Together these findings suggest LCs as key players in the pathophysiology of PDN in both mice and humans.

### Depletion of epidermal LCs prevented mechanical allodynia and spontaneous pain in the HFD model of PDN in male mice

To demonstrate that the LCs enrichment observed in the epidermis is relevant to PDN, we temporarily ablated LCs using a transgenic method while monitoring the development of mechanical allodynia and spontaneous pain in the HFD model of PDN. We conditionally expressed the Diphtheria Toxin Receptor (DTR) in LCs using a mouse line expressing DTR in langerin/Cd207 positive cells(37, 52). Weekly injections of Diphtheria Toxin (DT) starting at 4 weeks of HFD and continuing until 8 weeks of either RD or HFD temporarily ablated LCs in HFD DT-treated mice (**Fig. 3A)**. Histological analysis of paw epidermal sheets of HFD DT-ablated and control HFD (vehicle) showed the effective ablation of LCs after four DT injections. Indeed, the quantification of LCs density in epidermal sheets from male mice two days after DT injection demonstrated a 70% reduction of LCs density compared to mice injected with saline **(Fig. 3D-E).**

**Figure 3.**
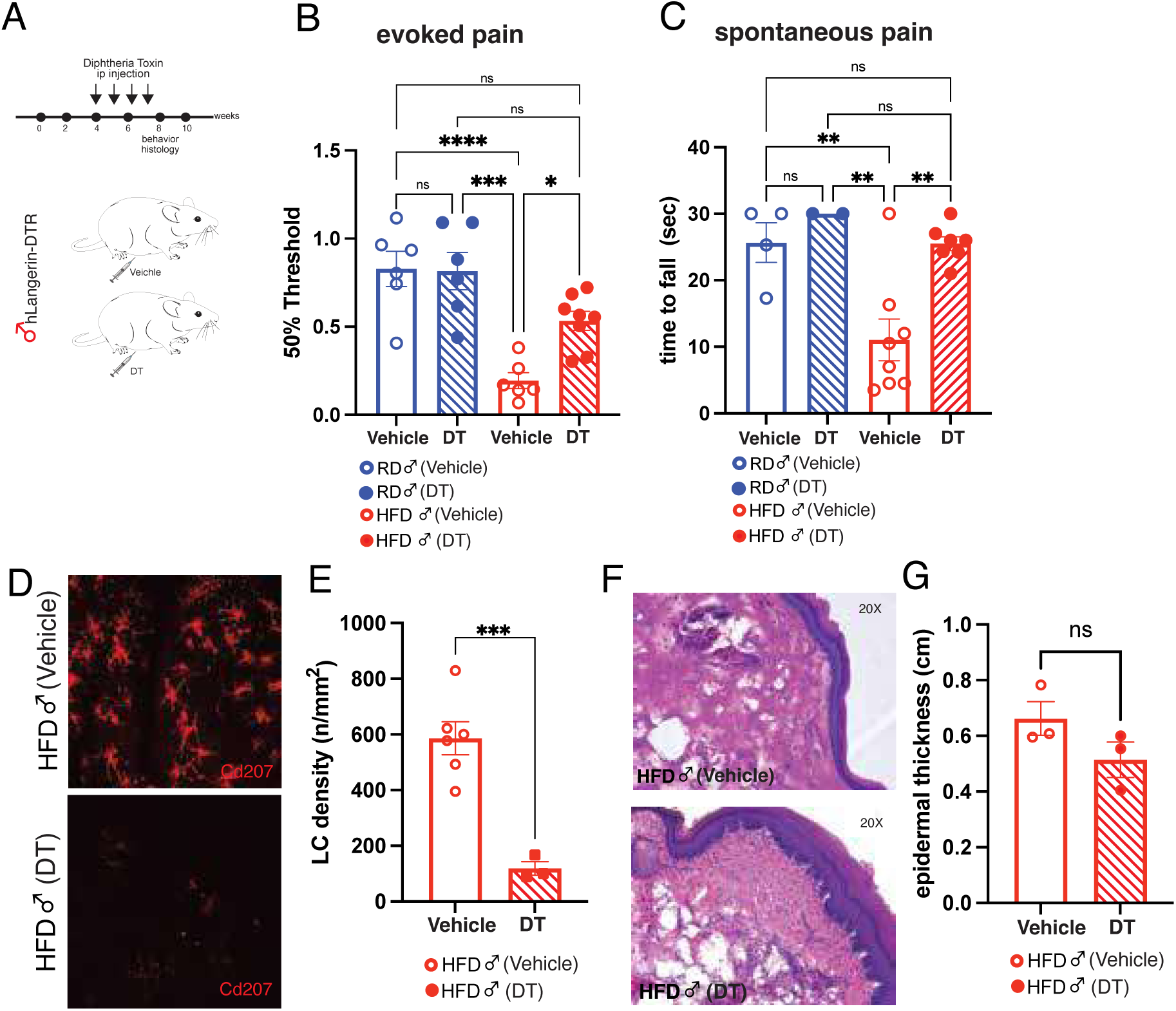
LCs mediate mechanical allodynia and spontaneous pain in HFD. **(A)** Timeline of diphtheria-toxin ablation strategy in hLC-DTR HFD male mice. **(B)** Response to evoked mechanical stimuli threshold. von Frey test shows the recovery of mechanical allodynia in HFD DT-ablated mice. One-way ANOVA for multiple comparison: RD (vehicle) vs HFD (vehicle) p<0.0001 (****); RD (vehicle) vs RD (DT) p=0.9995 (ns); RD (vehicle) vs HFD (DT) p=0.0514 (ns); HFD (vehicle) vs HFD (DT) p=0.0213 (*); HFD (vehicle) vs RD (DT) p=0.0001 (***); HFD (DT) vs RD (DT) p=0.0658 (ns). RD (vehicle) n= 6; RD (DT) n= 6; HFD (vehicle): n= 6 HFD (DT): n=8. **(C)** Spontaneous pain behavior measured as time to fall (seconds) is fully recovered by DT-ablation in HFD. One-way ANOVA for multiple comparison: RD (vehicle) vs HFD (vehicle) p=0.0084 (**); RD (vehicle) vs RD (DT) p=0.8627 (ns); RD (vehicle) vs HFD (DT) p>0.9999 (ns); HFD (vehicle) vs HFD (DT) p=0.0022 (**); HFD (vehicle) vs RD (DT) p=0.0081 (**); HFD (DT) vs RD (DT) p=0.8192 (ns). RD (vehicle) n= 4; RD (DT) n= 2; HFD (vehicle): n= 8 HFD (DT): n=7. **(D)** Representative confocal images of paw epidermal sheets of HFD male mice injected with vehicle (0.9% NaCl) or DT 4ng/gr body weight. **(E)** LCs’ density measured as number of Cd207+ cells/ area. Unpaired two-tailed t-Test with Welch’s correction p=0.0003 (***). HFD (vehicle): n= 4. HFD (DT): n=3. **(F)** Representative images of Hematoxylin-Eosin labeled skin sections of HFD male mice vehicle- and DT-injected. **(G)** quantification of epidermal thickness in HFD mice vehicle- and DT-injected. Unpaired two-tailed t-Test with Welch’s correction p=0.1681 (ns).

To determine if the presence of LCs in the epidermis is necessary and sufficient for the establishment of mechanical allodynia in HFD mice, we performed von Frey testing on both DT-treated and vehicle-treated RD and HFD male mice **(Fig. 3B).** Our results showed that the reduction of LCs through multiple DT injections during the HFD prevented the development of mechanical allodynia in the HFD male mice. In contrast, HFD mice that received vehicle treatment (0.9% NaCl) did not show any recovery from mechanical allodynia **(Fig. 3B**). Moreover, diphtheria toxin-induced LCs ablation did not affect the response to mechanical stimulation in RD DT-ablated male mice **(Fig. 3B).** Additionally, we measured spontaneous pain performing cage-lid hanging(53), and observed that HFD male mice developed robust spontaneous pain behavior compared to control RD **(Fig. 3C)** but this was recovered by DT-mediated LCs ablation **(Fig. 3C).** We confirmed that the DT-mediated strategy effectively ablated LCs HFD male mice **(Fig. 3D-E)**, without altering the histological organization of the epidermis **(Fig. 3F-G)**.

These results demonstrated that the presence of LCs in the epidermis is essential for maintaining mechanical allodynia and spontaneous pain in the HFD model of PDN in male mice. Next, we aimed to explore the mechanisms underlying this phenomenon.

### Single-cell transcriptional profiling of the epidermis in male mice reveals ten distinct clusters including LCs

To better understand the molecular mechanisms underlying the LCs-dependent development of neuropathic pain in the HFD mouse model of PDN, we elected to use an unbiased transcriptomic approach. The skin is a highly complex tissue; in addition to keratinocytes, the most abundant epidermal cell type, we observed other cell types including immune cells and LCs. To explore the complexity and heterogeneity of the epidermis in the well-established mouse model of PDN(39, 40), we performed scRNAseq of the paw epidermis of male mice fed an RD (RD n=3) and an HFD (HFD n=3) for 10 weeks **(Fig. 4A)**. After peeling the epidermis from the dermis, we obtained a high-viability single cell-suspension from each sample, collecting 40500 cells from RD epidermis and 28950 cells from HFD epidermis. We sequenced single cells using the 10x Genomics Chromium system. To remove doublets and low-quality cells, we filtered out cells with a number of features lower than 200 and higher than 6000 and containing more than 5% of mitochondria reads **(Suppl. Fig. 3A-G)**. After filtration, ten distinct clusters were identified based on the shared nearest neighbor (SNN) clustering algorithm in Seurat and visualized using the two-dimension Uniform Manifold Approximation and Projection (UMAP) method **(Fig. 4B).** To define all clusters, we integrated information of well-known epidermal markers with the top five differentially expressed gene markers of each cluster **(Fig. 4C),** identifying eight groups of keratinocytes at different stages of differentiation and two groups of non-keratinocytes. As the most abundant cell type in the epidermis, we used a combination of different markers to define the differentiation stages of keratinocytes **(Fig. 4D and Suppl. Fig. 3)**. Feature plots illustrated the distribution of keratinocytes markers, highlighting terminally differentiated keratinocytes, usually located in the upper layer of the epidermis corresponding to the *stratum corneum*, differentiated keratinocytes including spinous, suprabasal and squamous keratinocytes, undifferentiated basal epithelium keratinocytes mainly expressing Keratin14 (Krt14), migratory keratinocytes and keratinocytes with higher expression of Mki67,a distinctive marker of proliferation **(Fig. 4B-C).** We selectively identified: (1) terminally differentiated keratinocytes (tdKCs) expressing Flg, Krt78, Krt80, Lor and Ivl; (2) spinous keratinocytes (sKCs) expressing Cstdc5 and Dsg1a; (3–4) suprabasl keratinocytes type I (sbKCs I) and type II (sbKCs II) specified by Krt10 and Krt1, and Krt16, Krt17 and Il34, respectively; (5) squamous keratinocytes (sqKCs) expressing Krt16, Serpinb12 and Tmem266. To define undifferentiated KCs, we used (6) Krt14, Itga6 and Itgb1 for Basal Layer Keratinocytes (blKCs); (7) Krt79, Dcn, Lrig1 and Sema3e for migratory Keratinocytes (mKCs) and, lastly, (8) Mki67 and Cenpa for proliferating keratinocytes (pKCs) **(Fig. 4D-E)**. Furthermore, we used the high and selective expression of Cd207, Cd74, Cd28 and Cd3e to identify the remaining non-keratinocytes as (9) LCs and (10) immune cells, respectively **(Fig. 4D-E).** To validate some cluster markers in the paw skin sections of RD and HFD mice, we used keratin10 (Krt10) as marker of epithelial differentiation and Krt14 to label cells in contact with the basement membrane **(Fig. 4F-G).** We labelled epidermal LCs in paw epidermal sections of RD and HFD male mice using Cd207/Langerin (**Fig. 4H-I).**

**Figure 4.**
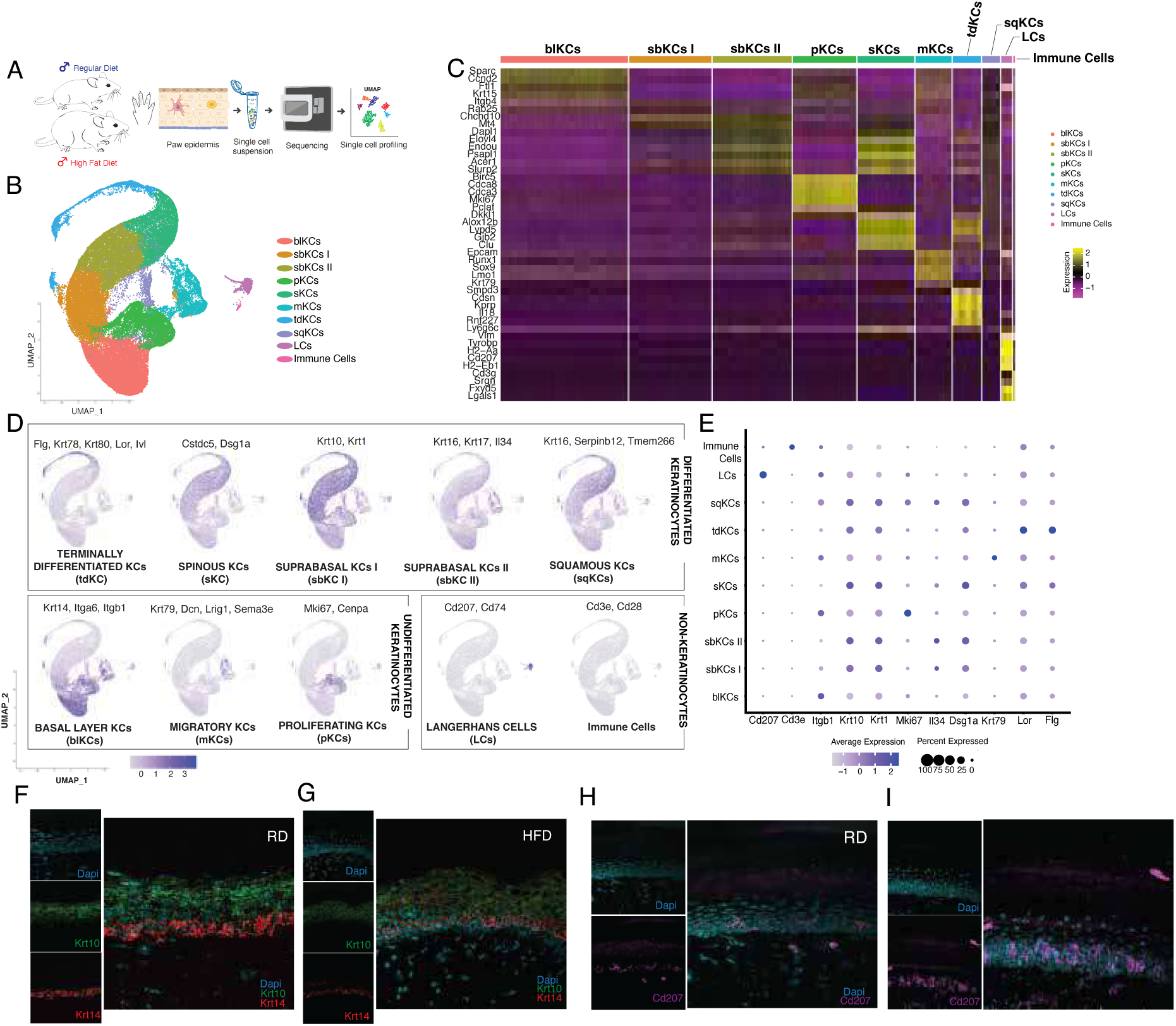
scRNAseq of paw epidermis of RD and HFD male mice. **(A)** Schematic workflow of scRNAseq of paw epidermis of RD and HFD male mice. **(B)** UMAP plot visualization of all 10 clusters identified in scRNAseq data of RD and HFD male paw epidermis. **(C)** Heatmap of top 5 differentially genes expressed in each cluster. **(D)** Feature plot map of each cluster showing the most representative gene markers. Terminally differentiated Keratinocytes (tdKCs) identified by Flg, Krt78, Krt80, Lor and Ivl. Spinous Keratinocytes (sKCs) identified by Cstdc5 and Dsg1a. Suprabasl keratinocytes type I (sbKCs I) identified by Krt10 and Krt1. Suprabasal Keratinocytes type II (sbKCs II) identified by Krt16, Krt17 and Il34. Squamous Keratinocytes (sqKCs) identified by Krt16, Serpinb12 and Tmem266. Basal Layer Keratinocytes (blKCs) identified nby Krt14 Itga6 and Itgb1. Migratory Keratinocytes (mKCs): Krt79, Dcn, Lrig1 and Sema3e. Proliferating Keratinocytes (pKCs): Mki67 and Cenpa. Non keratinocytes cells (1) LCs identified by Cd207 and Cd74; (2) Immune Cells identified by Cd3e and Cd28. **(E)** Dotplot visualization of the most representative markers of each cluster. The size of the dot describes the percentage of cells within a group (Percentage expressed) and the color indicates the Average Expression of each marker. **(F-G)** Representative confocal images of skin sections from RD and HFD paw male mice showing the expression and localization of undifferentiated Krt14+ (red) and differentiated Krt10+ (green) keratinocytes. **(H-I)** expression of LCs (Cd207/Langerin) in magenta in skin sections from paw male RD and HFD mice.

### Molecular and inflammatory profiling of LCs reveals differential release of inflammatory mediators in HFD model of PDN in male mice

To determine the molecular profile of various cell types in the epidermis related to PDN, we conducted a comparative clustering analysis of two dietary conditions. This analysis revealed differences in the diverse cell types present in the epidermis of mice on RD compared to those fed a HFD **(Fig. 5A**). We specifically focused on the LCs cluster because we have demonstrated that the presence of LCs in the epidermis is both necessary and sufficient for maintaining mechanical allodynia and spontaneous pain in the HFD mouse model of PDN (Fig. 1 and 4). Hence, to gain a deeper understanding of the molecular mechanisms involved in this process, we conducted Gene Set Enrichment Analysis (GSEA)(54) using the Mouse Molecular Signatures Database (MSigDB), which is one of the most comprehensive gene set databases available(55). Differentially expressed genes in RD and HFD LCs, identified using Seurat v5 (details in Methods), were analyzed using the GSEA package in R, and their expression profile was compared to MSigDB dataset. The significantly enriched gene sets were determined using the Enrichment Score corrected for multiple testing, obtaining the Normalized Enrichment Score (NES), where a positive NES value reflects the enrichment of gene sets on the ranked gene list, and, on the other hand, negative NES value indicates the opposite. We found that changes in the transcriptional profile of LCs between RD and HFD corresponded to an enrichment of genes involved in the MHC II-antigen presenting complex, contrary to the negative NES values mainly linked to synaptic protein translation or chemokine receptor binding **(Fig. 5B and Suppl. Fig. 4A)**. GSEA analysis suggested that the group of genes related to the MHC-II antigen presenting complex pathways was enriched in HFD **(Fig. 5C)**, contrary to gene sets for chemokine-receptors and synaptic proteins involved in translational mechanisms **(Fig 5D-E)**. To investigate potential molecules involved in any abnormal communication between LCs and sensory neurons, we combined information from the literature(30) and differentially expressed genes (DEGs) identified in LCs cluster **(Suppl. Fig. 4B).** Looking for genes linked to axonal guidance and immune responses, we detected the expression of Plxnb2 and Plxna1 in LCs **(Fig. 5f)** that was higher in HFD LCs **(Fig. 5G and Suppl. Fig. 4C-D),** as well as Itgam/Cd18, and H2-M2, involved in the immune signaling **(Fig. 5F-G)**.

**Figure 5.**
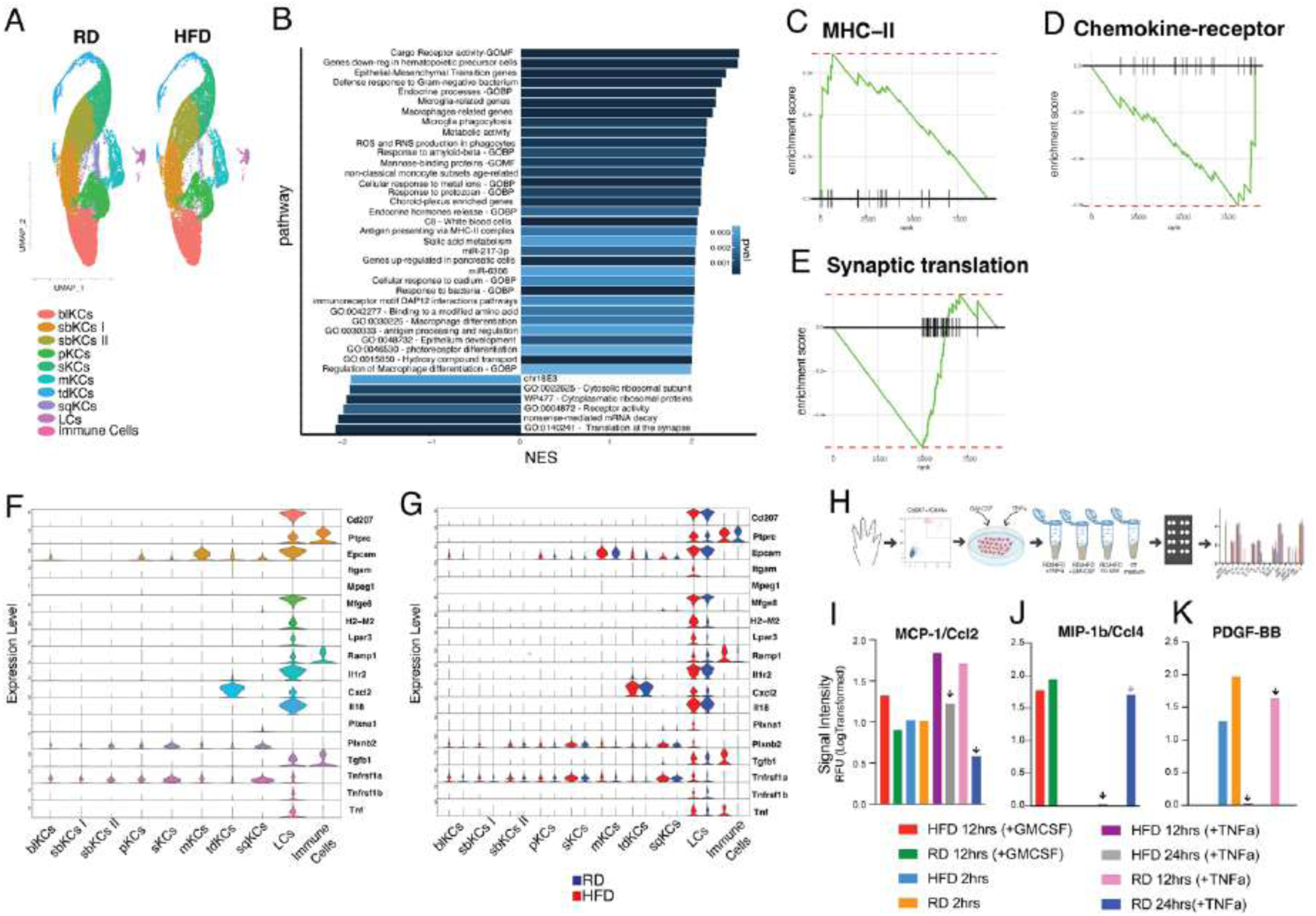
GSEA of LCs cluster. **(A)** UMAP plots of the scRNAseq of the two diet conditions, RD and HFD. **(B)** Plot shows GSEA of Langerhans cells cluster identifies enriched sets of genes with positive and negative NES value (abs(NES) > 1.88). **(C-E)** examples of major enriched pathways with positive NES showing the enrichment in MHC-II-related genes (c) and negative NES (d-e). Statistics were derived using the fgsea package in R. Adjusted p-value results from Benjamini-Hochberg correction. **(F-G)** Stacked violin plot showing the expression of genes implicated in axonal guidance, such as Plxna1 and Plxnb2, and inflammatory response. Combined (F) and split (G) by diet conditions gene expression (RD: blue – HFD: red). **(H)** Representative workflow of cytokines profile assay **(I-K**) Barplots show inflammatory molecules detected in the multiplexed proteomic profiling. Black arrows indicate differences for diet and treatment condition.

Given the known cellular and functional diversity of LCs(47, 56–58), we conducted a subcluster analysis of the major cluster of these cells. This analysis revealed four distinct molecular subclusters **(Suppl. Fig. 5A)**, each characterized by unique marker genes **(Suppl. Fig. 5A)**, which might be involved in different functions in PDN. We identified (1) Epcam+/Cd48+ LCs, (2) Ly6g6c+ LCs, (3) Cenpe+ LCs and (4) Ccr7+ LCs. As expected, all four clusters expressed the common Langerhans cells’ marker Cd207 **(Suppl. Fig. 5B-C)**. Interestingly, we found that only the subcluster 3 selectively expressed Ccr7, a marker for migrating LCs **(Suppl. Fig. 5B, D).** LCs are reported to express Ccr7 during the migration from the epidermis to the dermal layer to reach the lymph nodes and interact with T-cells(44, 59). The expression of Ly6g6c in two subclusters (cluster 1 and 3) indicated the presence of LCs associated with monocyte origins and it was increased in HFD LCs **(Suppl. Fig. 5B, E).** In HFD, LCs, similarly to resident macrophages, can recruit Ly6g6c+ cells from the blood contributing to the higher number of cells observed in the epidermis(56, 60). We also observed a broad expression of the membrane glycoprotein Epcam (Epithelial Cell Adhesion Molecule) **(Suppl. Fig. 5M, N)**, fundamental for the adhesion of LCs’ dendrites within epidermal keratinocytes(61). Interestingly, Cd48, even though expressed by all four clusters, was higher in HFD LCs **(Suppl. Fig. 5M, N)**. Cd48, a member of the signaling lymphocyte activation molecule family, is usually increased under inflammatory conditions and is widely expressed by immune cells(62), including LCs(46), where it could help in regulating immune responses through the activation of T-cells(46, 62). The selective expression of Cenpe (Centromere-associated protein E) in cluster 2 **(Suppl. Fig. 5M, N)** suggested that a small number of LCs might undergo local cell division and proliferation. Cenpe has been shown to be expressed by different immune cells, including macrophages and dendritic cells(63). We were also able to detect the expression of TNF-*α* and TGF-β1, a fundamental immune regulator for LCs activity and differentiation **(Suppl. Fig. 5P, Q)**. Moreover, subclustering analysis showed a broad expression of the chemokine Ccl22, usually expressed by LCs and dermal dendritic cells after injury(64, 65) **(Suppl. Fig. 5O)**. However, Ccl22 appeared downregulated in HFD, indicating a potential imbalance between these two subpopulations in HFD. Interestingly, the Lysophosphatidic acid receptor 3 (Lpar3) known to be a marker of non-peptidergic nociceptors type 1 (NP1)(39, 66, 67) was highly expressed in HFD LCs **(Suppl. Fig. 5F),** as well as Ramp1, co-receptor of calcitonin gene-related peptide receptor (CGRP)(68) **(Suppl. Fig. 5G)**,. Among several inflammatory molecules involved in the immune response mediated by LCs, we found that Interleukin-18 (Il18) was upregulated in HFD **(Suppl. Fig. 5H)**. We were also able to detect the transcript of both Mcp-1/Ccl2 and Mip-1b/Ccl4 in HFD LC subclusters **(Suppl. Fig. 5K-L)**, both molecules being involved in mediating neuropathic pain(69, 70).

LCs are key players in the innate immune response(25). When activated, these cells secrete pro-inflammatory cytokines and release nitric oxide, which might sensitize DRG axonal afferents in the epidermis(18). To reveal the unique secretome signature of LCs in the HFD model of PDN in male mice, we performed a multiplexed proteomic profiling of sorted Cd207+/Cd45+ LCs. A codeplex secretome analysis for innate immune response was able to detect the expression of different inflammatory molecules selectively secreted by LCs **(Fig. 5H)**. In addition to TNF-*α* and the granulocyte-macrophage colony-stimulating factor (GM-CSF), as expected, we detected the elevated expression of interleukin-10 (IL-10) and the macrophage migration inhibitory factor (MIF) **(Suppl. Fig. 5R-S).** We found that monocyte chemoattractant protein-1 (MCP-1/Ccl2) was secreted more by HFD LCs after TNF-*α* stimulation (24h after TNF-*α*) **(Fig. 5I)**, contrary to the macrophage inflammatory protein (Mip-1b/Ccl4) secreted only by RD LCs stimulated by TNF-*α* (24h) **(Fig. 5J).** Another interesting candidate differentially secreted by LCs is platelet-derived growth factor-bb (PDGF-BB), a potent mitogenic factor. After 12h of TNF-*α*, only RD LCs secreted PDGF-BB **(Fig. 5K)** suggesting that the synergistic action between TNF-*α* and PDGF-BB is potentially deregulated in HFD. The known role of Mcp1/Ccl2 in neuropathic pain(69, 70) strongly suggests the inflammatory activity mediated by LCs contributes to maintaining mechanical allodynia in HFD.

### Altered neuron-immune communication between LCs and cutaneous afferents via Semaphorin-Plexin signaling pathways in PDN

In the epidermis there is significant interaction between cutaneous afferents and non-neuronal cells, including keratinocytes and LCs^4^. To understand how LCs coordinate interactions with other cells, especially with nerve afferents in the epidermis, we adopted CellChat, an open source R package (https://github.com/sqjin/CellChat) capable of disentangling intricate signaling networks(71). By using paw epidermis scRNAseq data from male mice, CellChat predicted potential cell-cell communication based on the over expressed ligands and receptors identified in each cell group within the epidermis **(Fig. 6A).** We observed that the number of inferred examples of cell-cell communication, as well as the strength of cell-cell interactions, was increased in HFD **(Fig. 6B-C and Suppl. Fig. 6A).** We also compared the information flow for all signaling pathways, defined by “*the sum of communication probability among all pairs of cell groups in the inferred network*” **(Suppl. Fig. 6B-C)**. Heatmap visualization allowed us to obtain an overall view of all identified pathways and their relative strength in each cluster in RD and HFD, highlighting dramatic differences in molecular signaling pathways between RD and HFD **(Fig. 6d).** We observed that among keratinocytes clusters, proliferating (pKCs) and terminally differentiated (tdKCs) keratinocytes are extremely influenced by changes in signaling pathways such as the pleiotrophin (PTN) or Epidermal Growth Factor (EGF) pathways, which are also found to be upregulated during wound repair and healing processes(72), or cell adhesion molecules (Nectin). Examining potential signaling pathways involved in neuronal contact and communication, we found that Sema6 and Sema4 were differentially modulated in HFD compared to RD **(Fig. 6D).** The versatile role of semaphorins in regulating cell shape and motility during the development, axonal guidance and immune responses, make them good candidates for investigating communication between LCs and nerve afferents in the epidermis. Cell-cell communication analysis suggested robust changes in Sema4 and Sema6 signaling pathways. For example, in examining the Semaphorin 4 (Sema4) pathway, we found that LCs do not interact with any other cell types in the epidermis under RD conditions **(Fig. 6E)**. However, during HFD, LCs become targeted by proliferating squamous keratinocytes **(Fig. 6E).** Additionally, we observed that HFD LCs communicate with one another through an autocrine mechanism facilitated by Semaphorin 6 (Sema6) signaling **(Fig. 6F).** This analysis reveals that the Sema4 and Sema6 signaling pathways are differentially regulated in HFD LCs. Upon examining members of the Sema4 family, we found that only Sema4a is expressed in keratinocytes under both RD and HFD conditions. Similarly, only the candidate receptors Plxnb1 and Plxnb2 were detected in epidermal cells, including LCs **(Suppl. Fig. 6D).** In contrast, our analysis of the Sema6 family revealed a significantly higher expression of Sema6d and its receptor Plxna1 in HFD LCs. This suggests that HFD LCs may have enhanced self-maintenance and self-activity through Sema6d-Plxna1 signaling (**Fig. 6F and Suppl. Fig. 6D).** However, intercellular communication within the epidermis alone cannot adequately explain the nerve damage observed in PDN.

**Figure 6.**
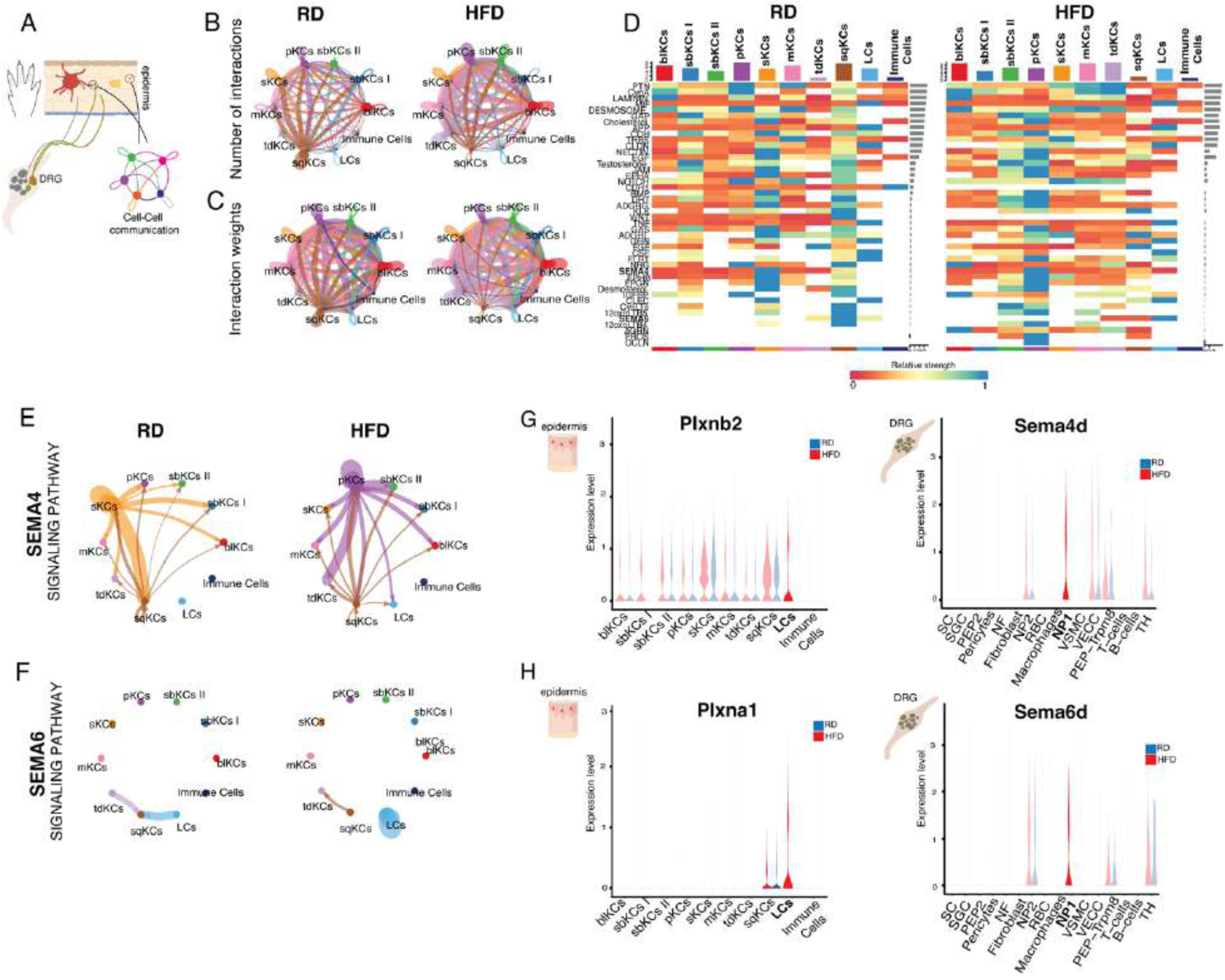
cell-cell communication between LCs and DRG sensory neurons. **(A)** Schematic cartoon of cell–cell ligand–receptor network analysis performed using CellChat. **(B-C)** Circle plots show the number of interactions (b) and the strength of interactions (c) between different cell types in RD/upper panels and HFD/lower panels. Circle sizes are proportional to the size of different cell types and the thickness of the lines indicates a stronger signal. **(D)** Comparison of cell-cell communication between RD and HFD using Heatmap plot. Highlighted in bold Sema4 and Sema6 signaling pathways. **(E-F)** Circle plot representing Sema4 and Sema6 signaling pathways, respectively, in RD and HFD. **(G-H)** Violin plots of expression levels of Plxna1 and Plxnb2 in the scRNAseq of paw epidermis (g) and Sema4d and Sema6d expression in scRNAseq of DRG (h) of RD and HFD male mice.

Various neuronal subpopulations terminate in the epidermis(6, 7), forming both gap junctions and synapse-like contacts with epidermal non-neuronal cells(7, 10–12, 73). This creates opportunities for communication between non-neuronal cells and nerve afferents. To investigate how the interactions between LCs and nerve afferents in the epidermis might be affected in PDN, we employed a combined approach utilizing scRNAseq data from the epidermis along with previously published scRNAseq data from the DRG of mice fed a RD or a HFD(39) **(Fig. 6A).** We specifically examined the expression of members from the Sema4 and Sema6 families. Notably, in our analysis of the scRNA seq data from DRG neurons, we found that Sema6d is present only in the non-peptidergic nociceptor type 1 (NP1) subtype of DRG neurons in HFD mice. Additionally, in the epidermis scRNA data set, the Sema6d ligand, Plxna1, is expressed exclusively in LCs from HFD mice **(Fig. 6H and Supp. Fig. 6E).** Likewise, Sema4d is expressed in NP1 neurons from HFD mice, and its ligand, Plxnb2, is found in LCs from HFD mice **(Fig. 6H and Supp. Fig. 6E).** To validate our findings, we conducted in-situ RNA scope analysis on frozen sections of DRG from mice fed either a RD or HFD. In these validation studies, we confirmed that Mrgprd-positive neurons, identified as NP1, also express Sema4d **(Suppl. Fig. 6F-G)** and Sema6d **(Suppl. Fig. 6H-I)**. NP1 afferents, also known as Mrgprd-positive afferents, play a crucial role in attracting LCs in the skin(37). Additionally, the survival of NP1 afferents is linked to their association with these LCs(37). Together, these findings suggest that altered communication between neurons and the immune system, specifically between LCs and NP1 Mrgprd-positive cutaneous afferents, may involve semaphorin-plexin signaling pathways. This altered communication could contribute to the remodeling of cutaneous innervation in PDN.

### Dimorphic molecular signatures of LCs in the HFD model of PDN

Several indications suggest that sex differences are a fundamental biological variable influencing immune cells, particularly microglia(74–76) and, consequently, immune responses(77) and neuropathic pain(18, 78). Therefore, we decided to investigate the role of LCs in PDN in HFD female mice. We first characterized the HFD phenotype in female mice after 10 weeks of diet **(Suppl. Fig. 7A-B)**. Even though the 42% of fat diet led to a robust increase of the body weight in all female subjects (**Suppl. Fig. 7A**), only in a few did the blood glucose level reached the cutoff 120 minutes after glucose injection (additional information in Methods) (**Suppl. Fig. 7B**). Thus, we measured the response to mechanical forces only in high-glucose level females determining the onset of mechanical allodynia by quantifying the withdrawal threshold of the hind-paw in response to von Frey stimulation. We found that, similarly to male mice(39, 40), female mice fed a high-fat diet developed mechanical allodynia after 10 weeks (**Suppl. Fig. 7C**).

In the HFD mouse model of PDN, we observed a significant increase in LCs in the whole-mount epidermis of wild-type male mice that were fed a HFD for 10 weeks, and this increase was not seen in male mice on RD (Fig. 1a-b). To determine if similar results occurred in HFD female mice, we quantified LC density in the epidermis using epidermal sheets from both RD and HFD female mice. Interestingly, we found that HFD did not lead to an increase in LC density in the paw epidermis of female mice **(Fig. 7A-B).** Furthermore, when comparing the density of LCs in male and female mice on a RD and a HFD, we observed that female RD mice had a higher number of LCs than male RD mice **(Suppl. Fig. 7D).** This significant difference between male and female mice suggests that the role of LCs in PDN may be influenced by sex.

**Figure 7.**
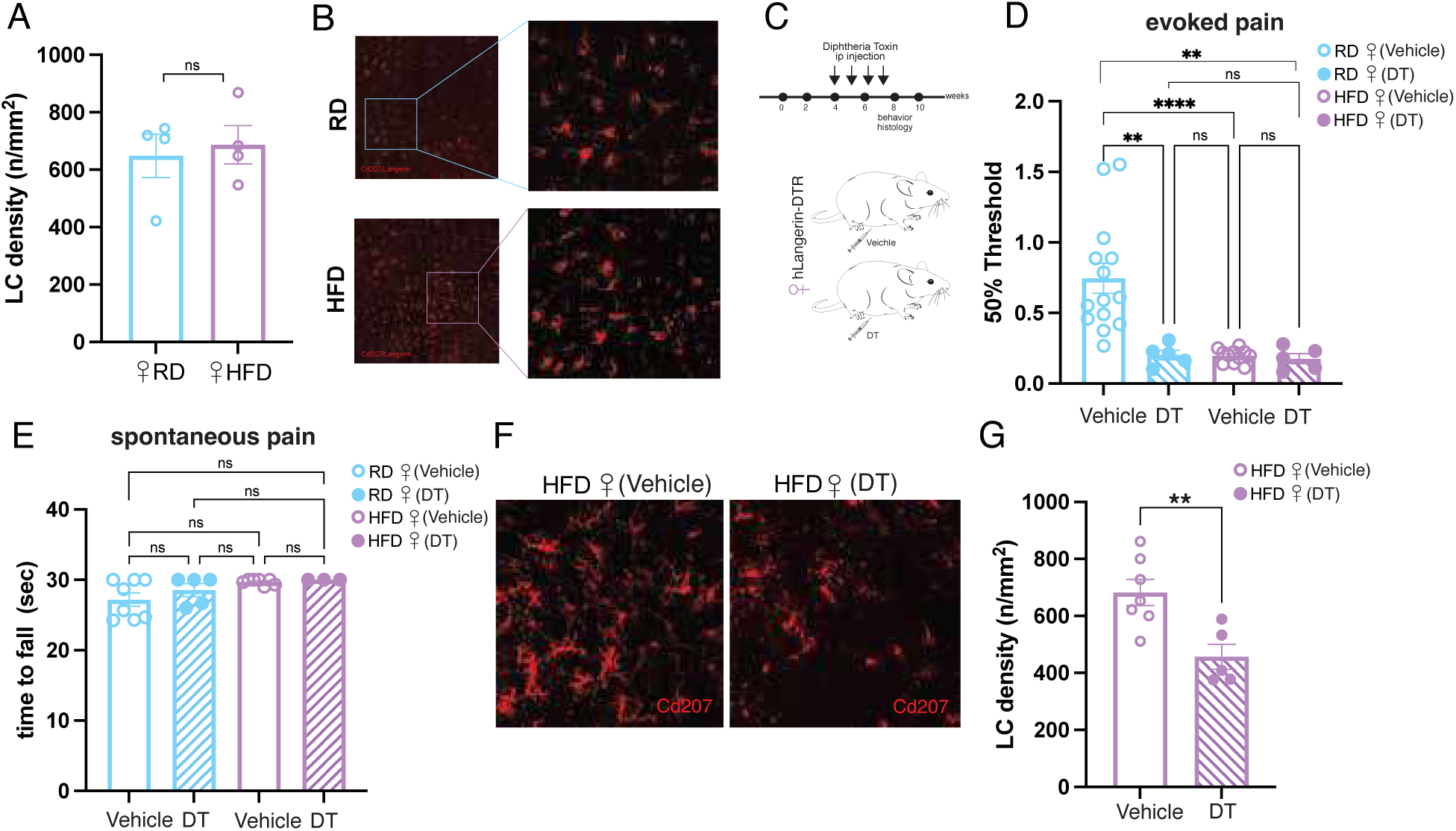
LCs involvement in female HFD. **(A-B)** Quantification of LCs’ density (number of LCs / mm^2^) in RD and HFD female mice at 10 weeks (b) and representative images (c). n=3 sections per animal were acquired and Cd207+ cells per 0.04 mm^2^ per section area were counted. Unpaired two-tailed t-Test p=0.7154. n=4 for both groups. **(C)** schematic timeline of DT-mediated ablation strategy in HFD female mice. **(D)** Response to evoked mechanical stimuli threshold through von Frey test. DT-ablation does not recover mechanical allodynia in HFD female. One-way ANOVA followed by Tukey’s multiple comparison. RD (vehicle) vs RD (DT) p=0.0022 (**), RD (vehicle) vs HFD (vehicle) p<0.0001 (****), RD (vehicle) vs HFD (DT) p=0.0013 (**). RD (vehicle): n= 14, RD (DT): n=5, HFD (vehicle) n=11; HFD (DT) n=5. **(E)** Spontaneous pain behavior measured as time to fall (seconds) is not affected by DT-ablation in HFD female. One-way ANOVA followed by Tukey’s multiple comparison. RD (vehicle): n= 8, RD (DT): n=5, HFD (vehicle) n=7; HFD (DT) n=3 **(F)** Representative confocal images of paw epidermal sheets of HFD female mice injected with vehicle (0.9% NaCl) or DT 4ng/gr body weight. **(G)** LCs’ density measured as number of Cd207+ cells/ area. Unpaired two-tailed t-Test with Welch’s correction p=0.0055(**). HFD (vehicle): n= 7. HFD (DT): n=5.

To further explore the role of LCs in female mice with PDN, we temporarily ablated the LCs using the same transgenic method employed in male mice **(Fig. 7C)**. First, we conditionally expressed the Diphtheria Toxin Receptor (DTR) in LCs by utilizing a mouse line that expresses DTR in langerin/CD207-positive cells(37). We then monitored the development of mechanical allodynia and spontaneous pain in the HFD model of PDN. While multiple injections of DT robustly ablated LCs in male mice (Fig. 4d-e) without affecting the anatomy of the epidermis (Fig. 4f-g), the same ablation strategy used in male mice resulted in only a 40% of reduction of LCs in female HFD DT-ablated mice **(Fig. 7F-G).** Interestingly, LCs ablation in HFD female mice did not prevent mechanical allodynia but, instead, it was responsible of eliciting pain behavior in RD DT-ablated mice **(Fig. 7D).** This suggested that pain mechanisms mediated by LCs differ between male and female mice. Moreover, contrary to male HFD mice, we found that female HFD mice did not develop spontaneous pain and LC ablation DT-mediated does not affect this pain behavior in female mice, both in HFD and RD **(Fig. 7E).**

To better understand the molecular mechanisms underlying the possible dimorphic role of LCs in this PDN model, we extended the single-cell transcriptomic analysis to the epidermis of female mice in the HFD model of PDN. We performed scRNAseq of the paw epidermis of female mice fed a regular diet (RD n=3) and a high-fat diet (HFD n=3) for 10 weeks **(Fig. 8A)**. After peeling the epidermis from the dermis, we obtained a high-viability single cell-suspension from each sample, collecting 33175 cells from RD epidermis and 31047 cells from HFD epidermis. We applied cell filtering based on quality control metrics, retaining cells with several detected features (nFeature_RNA) greater than 500 and less than 6000, and the mitochondrial gene content (percent.mt) below 5%. This filtering step helped improving data reliability by excluding low-quality or dying cells **(Supp. Fig. 8A-B).** Similarly to male scRNAseq data (Fig. 5), we identified 10 molecularly distinct clusters mainly represented by keratinocytes but also immune cells and LCs **(Fig. 8B**). For each cluster, we analyzed the top 10 differentially expressed genes, revealing key gene signatures that characterized each cell population **(Fig. 8C)**. To determine sex-mediated differential expression of specific target genes linked to axonal guidance and immune responses, we analyzed a panel of genes detecting the higher expression of Plxna1 and a lower expression of Plxnb2 in LCs **(Fig. 8D, G-H)**. We also performed a sub-clustering analysis of the LC cluster in the female scRNAseq dataset identifying four molecularly distinct clusters such as (1) Epcam+/Cd48+ LCs, (2) Ly6g6c+ LCs, (3) Ccr7+ LCs and (4) Cenpe+ LCs **(Fig. 8E-F).** To identify transcriptional changes in LC subclusters between RD and HFD, we applied a cutoff value of a P-value (adjusted for Benjamini-Hochberg correction) ≤0.05 and an absolute log_2_ fold change of 0.25 **(Supp. Fig. 8C).** We observed that the sodium voltage-gated channel alpha subunit 3 (Scn3a) and purinergic receptor P2X2 (P2xr2) were upregulated in female HFD LCs **(Supp. Fig. 8C).** Interestingly, by comparing the expression of Scn3a and P2xr2 in female and male LCs subclusters **(Fig. 8I-K),** we found that Scn3a was abundantly upregulated in female LCs **(Fig. 8I)** but absent in male LCs **(Fig. 8K)** and no differences were detected in male P2xr2 expression. The studies comparing LCs in female and male mice indicate that LCs may have distinct dimorphic functions as immune cells driving disease in PDN.

**Figure 8.**
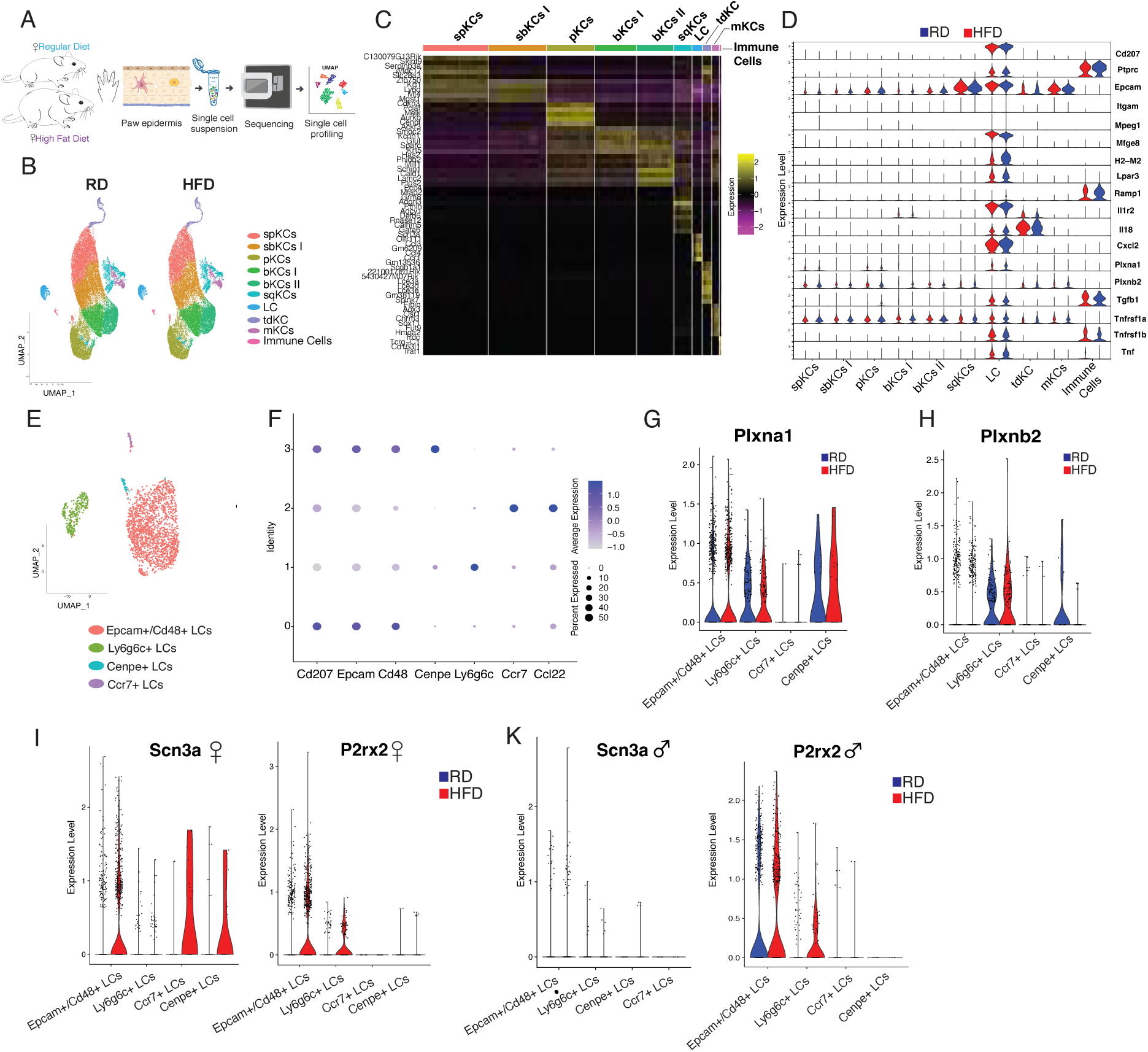
Transcriptomic profile of LCs in RD and HFD female mice. **(A)** Schematic workflow of scRNAseq of paw epidermis of RD and HFD female mice. **(B)** UMAP dimensionality reduction of RD and HFD scRNAseq paw epidermis of female mice shows ten distinct clusters identified by the expression of differentially expressed genes (DEGs). **(C)** Heatmap of top10 DEGs expressed in each cluster. **(D)** Stacked violin plot showing the expression of genes implicated in axonal guidance and inflammatory response split by diet conditions identified in the male scRNAseq data set (Fig. 5g). RD: blue – HFD: red. **(E)** UMAP dimensionality reductions of female LCs subclustering. Four distinct clusters were identified. **(F)** Dotplot shows the expression of genes adopted to identify LCs’ subclusters. **(G-H)** expression of Plxna1 (g) and Plxnb2 in LCs’ subclusters. (**i)** Violin plots show the expression of two of the DEGs upregulated in LCs female: Scn3a (left) and P2rx2 (right). **(K)** Expression of Scn3a (left) and P2rx2 (right) in male LCs subcluster.

## DISCUSSION

In these studies, we revealed, for the first time, a dimorphic role for LCs in the development of mechanical allodynia and spontaneous pain in the HFD model of PDN. We demonstrate that in PDN, the density of LCs increased in the paw epidermis of HFD-fed male mice correlating with mechanical allodynia and sensory fibers loss. Importantly, we found that LCs increase is linked to a sensory nerve degeneration in skin biopsies from well-characterized PDN patients compared to healthy controls. Crucially, we discovered that selectively ablating LCs using a diphtheria-toxin strategy prevented nociceptive behavior in male mice. Interestingly, LCs ablation in HFD female mice did not prevent mechanical allodynia but, instead, it led to pain behavior in DT-ablated female mice fed a RD. This suggested that pain mechanisms mediated by LCs differ between male and female mice. scRNAseq of the paw epidermis from both male and female HFD mice identified sex-mediated differences in the expression of specific target genes within this PDN model. In male mice, scRNAseq of LCs revealed differentially expressed genes related to immune responses and axonal guidance, including those involved in Semaphorin-Plexin signaling pathways. Furthermore, through a high-multiplex cytokine profiling assay, we identified a panel of inflammatory molecules secreted by LCs that may contribute to the onset and maintenance of PDN, such as Mcp1/Ccl2, which was highly secreted by LCs from HFD male mice. These findings highlight the role of LCs as key contributors to PDN and suggest their potential as therapeutic targets for innovative interventions, particularly topical treatments that modulate immune cell activity and neuroimmune communication in the skin. Moreover, the differences observed in female and male mice indicate that LCs may have distinct dimorphic functions as immune cells driving disease in PDN.

LCs, which make up about 2% to 5% of all epidermal cells(51, 79), are primarily known for their role in tissue surveillance, extending and retracting their cellular processes between keratinocytes(80, 81). Located in the outermost layer of the skin, LCs are crucial not only for monitoring the epidermis but also for regulating innate immunity and recruiting adaptive immune T cells(25). The diverse range of cytokines they release in both physiological(30) and pathological(20, 82, 83) conditions add complexity to our understanding of the specific roles of individual inflammatory molecules. By employing a high-multiplex cytokine profiling assay, we identified various immune mediators that are selectively released by LCs and highlighted differences in their secretion between HFD and RD, including the expression of chemokine monocyte chemoattractant protein-1 Mcp-1/Ccl2 (Fig 5). Mcp-1/Ccl2, acting through its chemokine receptor Ccr2, has been reported to promote pain hypersensitivity in naïve rodents(84–87). Neuropathic pain caused by nerve injury(86) or chronic compression(88) leads to a significant upregulation of the Ccl2-Ccr2 signaling complex at both the nerve site and DRG cell bodies, an effect that is fully reversed in Ccr2-deficient mice. Moreover, Mcp-1/Ccl2 application has been reported to mediate DRG hyperexcitability through Ccr2-activation(88–90). Consistent with the pro-nociceptive function of Mcp-1/Ccl2, we found that HFD LCs secreted higher levels of Mcp-1/Ccl2 compared to RD LCs. However, the direct impact of Mcp-1/Ccl2 on DRG excitability in the HFD mouse model of PDN requires further investigation. Interestingly, in addition to this pro-nociceptive function of Ccl2/Ccr2 complex, recent evidence in diabetic neuropathy suggests that Ccr2-expressing macrophages can infiltrate damaged tissue to promote pain resolution(91).

scRNAseq applied to HFD paw epidermis enabled the exploration of the transcriptional signature of LCs, particularly their broader cytokine expression profile (Fig. 5 and Suppl. Fig. 4 and 5). LCs mobilization, which usually involves between the 20% and 30% of resident LCs, is a critical process following tissue damage and relies on the release of specific cytokines(92). Interleukin-18 (Il-18), a member of the Il-1 superfamily structurally similar to Il-1β, plays a key role in LCs mobilization process(92) and pro-nociceptive activity, as observed in animal models of neuropathic pain caused by nerve ligation(93) or chronic constriction injury(94). Notably, administration of Il-18 binding protein (Il-18bp), attenuates neuropathic pain by reducing Il-18 levels(94). Interestingly, the upregulation of Il-18 is not confined to neuropathic pain caused by traumatic injuries. Elevated levels of this cytokine have also been detected in the serum of individuals with type 2 diabetes mellitus(95) and pre-diabetes syndrome(96) compared to healthy controls. Our analysis indicates an upregulation of the Il-18 transcriptome in HFD LCs, which aligns with findings reported in the literature(96). Like the role of spinal microglia in neuropathic pain conditions(97), HFD LCs may contribute to sustained pain hypersensitivity in HFD mice through the release of elevated level of Il-18 in the peripheral environment of the epidermis.

As observed in both PDN patients(3, 4, 34) and animal models, such as HFD mice(39, 40), there is a massive degeneration of sensory fibers, particularly those innervating the epidermis. Following degeneration, physiological regeneration has been observed in skin biopsies of healthy subjects(98), as well as in mice with neuropathic pain, where aberrant end-organ reinnervation(99) or axonal sprouting(84, 100) occurs. Through unbiased transcriptomic analysis, we identified Semaphorin-Plexin signaling pathways that underlie the interaction between LCs and sensory neurons in HFD conditions. Semaphorins (Sema), which are membrane-bound or soluble molecules, along with their transmembrane receptors, Plexins (Plxn), are key modulators of axonal guidance in the developing nervous system(101, 102), as either attractants or repellents for neurites(103). While transcriptomic analysis does not fully elucidate the molecular mechanisms and does not provide direct evidence of the role of Sema-Plxn in modulating DRG neurite degeneration or regeneration, accumulating evidence suggests that NP1 Mrgprd-positive sensory neurons and LCs engage in mutual interactions(37). Indeed, from previous work we know that Mrgprd-positive afferents control the recruitment of LCs in the skin(37). Importantly, the survival of Mrgprd-positive afferents depends on their association with LCs(37). Beyond their role in guiding axons, Sema-Plxn signaling pathways have also been implicated in mediating immune responses(104–107) and in promoting nociceptive behavior in cases of pathological pain(108). Interestingly, semaphorins, through the activation of plexins, promote the activation of dendritic cells and their migration to lymph nodes(109, 110). Furthermore, Sema-Plxn signaling pathways promote the polarization of macrophages toward an anti-inflammatory state(106), suggesting a role in the regulation of antigen-presenting cells. Consistent with our findings, Plxnb2, which binds class IV transmembrane semaphorin ligands including Sema4d, is upregulated in microglia and macrophages after nerve injury(111). Blocking Sema-Plxn signaling has been shown to impair the development and duration of pain in a mouse model of inflammatory pain(108). However, further experiments using conditional Plxnb2 knockout transgenic mice are needed to prove the functional relevance of Sema-Plxn pathways in modulating DRG neurite degeneration or regeneration in PDN. Recent findings indicate sex-dependent mechanisms adopted by immune cells, especially resident macrophages such as microglia(74–76) and LCs, in various diseases. Similar to our observations on LCs in PDN male mice, studies using neuropathic pain animal models caused by nerve injury have shown the proliferation of spinal microglia exclusively in male animals(112). This suggests that LCs may also exhibit sex-specific behavior in PDN, as previously found for microglia in obesity(113). Notably, while LCs ablation did not affect pain behavior in RD DT-ablated male mice, our data on RD DT-ablated female mice suggest that an intact LCs network in the epidermis is necessary for pain sensation, and the loss of LCs in female rodents can trigger mechanical allodynia in naïve condition (Fig. 7). Moreover, transcriptomic analysis further demonstrated intrinsic sex differences in LCs gene expression, potentially linked to distinct activity and function in male and female rodents. We observed that the Scn3a and P2xr2 were upregulated in female HFD LCs (Fig. 8). Interestingly, by comparing the expression of Scn3a and P2xr2 in female and male LCs subclusters (Fig. 8), we found that Scn3a was abundantly upregulated in female LCs but absent in male LCs (Fig. 8). Voltage-gated sodium channels, including Scn3a, have been observed in several non-excitable cells, including macrophages and dendritic cells(114), but their role is not fully understood. Ion channels investigated in monocytes and microglia might mediate several cellular processes in activated cells such as proliferation, migration and neurite ramification in a sex-dependent way(115) (74–76). Sex-specific differences in the transcriptional profile of LCs between male and female mice might explain the differential outcomes observed in females after DT-mediated ablation of LCs (Fig. 4 and 10). For instance, the selective presence of Scn3a, only in female HFD LCs may influence their sensitivity to environmental inflammatory modulators and regulate the release of cytokines, as previously found in dendritic cells(116). However, a more comprehensive comparison of the LCs transcriptome of males and females, first in naïve RD and under HFD conditions, is an important aim for future study. The possible dimorphic role of LCs in mechanical sensation under normal physiological circumstances and in pathological conditions such as PDN is highly relevant to the study of peripheral neuropathic pain. If these findings are confirmed in human studies, they would highlight the importance of considering sex differences in developing more effective treatments. Thus, future research will concentrate on addressing this intriguing question.

To enhance the translational relevance of our studies, we analyzed key parameters from skin biopsies obtained from a cohort of clinically well-characterized patients with PDN and healthy controls (Fig. 2). Although the density of LCs remained unchanged in the skin sections of PDN patients (Suppl. Fig. 2), we discovered that the ratio of fiber denervation to LCs density was significantly reduced in PDN patients (Fig. 2). This finding suggests that the loss of cutaneous innervation in the epidermis of PDN patients is related to the density of epidermal LCs. Indeed, since LCs morphology is linked to their function(117), our observation of an increase in the volume and complexity of LCs arborization as the duration of diabetes increases in PDN patients (Fig. 2) may indicate potential structural and functional adaptations or compensatory mechanisms within LCs in the epidermis of PDN patients. We acknowledge the limitations of conducting these studies in humans, as PDN patients exhibit a variety of symptoms and sensory phenotypes(117). Therefore, there is a critical need for studies involving a larger cohort of clinically characterized PDN patients to explore the specific role of LCs in different sensory phenotypes.

In conclusion, these studies increase our knowledge of the interactions between non-neuronal cells and nerve endings in the epidermis and highlight the role of LCs in PDN by integrating data from the HFD animal model and human observations. We found that the increased density of epidermal LCs contributes to mechanical allodynia and spontaneous pain in male mice fed an HFD. A disrupted functional relationship between LCs and sensory afferent neurons in the epidermis may lead to the degeneration of sensory fibers, a characteristic feature of PDN seen in both patients and HFD mice. Recognizing LCs as immune cells that drive disease development suggests possibilities for creating topical therapeutic strategies targeting these cells in the epidermis. Potential approaches could include antigen-specific immunotherapies. LCs are a distinct and well-defined population of immune cells located in the upper layers of the epidermis. Their unique morphology makes them suitable candidates for intraepithelial treatment administration or for delivering therapies aimed at modulating their density and cytokine production in PDN. Furthermore, our studies in both male and female mice suggest that LCs may play different roles in mechanical sensation under both normal and pathological conditions, underscoring the importance of considering sex differences when developing more effective treatments for patient suffering from PDN.

## METHODS

### Study Design

This work describes the dimorphic role for LCs in the development of mechanical allodynia and spontaneous pain in a HFD mouse model of PDN. We utilize a variety of approaches, including transgenic methods, pain behavioral assessments, and single-cell transcriptomics in a clinically relevant HFD mouse model of PDN and skin samples from well-clinically characterized patients with PDN. This research complies with all relevant ethical regulations. All animal care protocols and experiments were reviewed and approved by the Institutional Animal Care and Use Committee (IACUC) of Northwestern University. Human tissues were collected from healthy volunteers and patients and the study was approved by the local Ethical Committee of the Fondazione IRCCS Istituto Neurologico ‘Carlo Besta’ of Milan (FINCB), Italy.

All procedures were designed to maximize robustness and minimize bias. Experimenters performed and analyzed LCs density in whole-mount epidermal sheets, RNAscope and behavioral assays were blinded to diet treatment (RD or HFD) and/or compound (diphtheria toxin or vehicle) until data analysis is complete. Both male and female mice were included in all experiments and results were analyzed separately and compared. To collect human tissues from PDN patients and controls, we first performed clinical examination of all subjects and obtained informed consent to participate in the study. For the present study, 15 patients with diabetic neuropathy (7 females and 8 males) and 9 healthy controls (4 females and 5 males) were enrolled. Clinical features were collected using an established protocol previously described(118). Pain intensity was measured as the average score of the last 3 weeks using the pain intensity numerical rating scale (PI-NRS).

Further information as number of animals used, replications, number of sections counted are indicated in the figures and/or figure legends. On the graphs, individual dots represent individual samples/mice used.

## Supporting information

Suppl. Materials includes extended methods sections and supplementary figures

## SUPPLEMENTARY MATERIALS

The file includes: (1) Detailed sections of Methods; (2) Statistics; (3) Data Availability; (4) Acknowledgements; (5) Fundings and (6) Supplementary Figures and legends.

## REFERNCES

1. Spallone V, Lacerenza M, Rossi A, Sicuteri R, Marchettini P. Painful diabetic polyneuropathy: approach to diagnosis and management. The Clinical journal of pain. 2012;28(8):726–43.

2. Dyck PJ, Kratz KM, Lehman KA, Karnes JL, Melton LJ, 3rd, O’Brien PC, et al. The Rochester Diabetic Neuropathy Study: design, criteria for types of neuropathy, selection bias, and reproducibility of neuropathic tests. Neurology. 1991;41(6):799–807.

3. Devigili G, Tugnoli V, Penza P, Camozzi F, Lombardi R, Melli G, et al. The diagnostic criteria for small fibre neuropathy: from symptoms to neuropathology. Brain. 2008;131(Pt 7):1912–25.

4. Lauria G, Devigili G. Skin biopsy as a diagnostic tool in peripheral neuropathy. Nat Clin Pract Neurol. 2007;3(10):546–57.

5. Sommer C, Lauria G. Skin biopsy in the management of peripheral neuropathy. Lancet Neurol. 2007;6(7):632–42.

6. Zylka MJ, Rice FL, Anderson DJ. Topographically distinct epidermal nociceptive circuits revealed by axonal tracers targeted to Mrgprd. Neuron. 2005;45(1):17–25.

7. Lumpkin EA, Caterina MJ. Mechanisms of sensory transduction in the skin. Nature. 2007;445(7130):858–65.

8. Sumner CJ, Sheth S, Griffin JW, Cornblath DR, Polydefkis M. The spectrum of neuropathy in diabetes and impaired glucose tolerance. Neurology. 2003;60(1):108–11.

9. Divisova S, Vlckova E, Srotova I, Kincova S, Skorna M, Dusek L, et al. Intraepidermal nerve-fibre density as a biomarker of the course of neuropathy in patients with Type 2 diabetes mellitus. Diabet Med. 2016;33(5):650–4.

10. Handler A, Ginty DD. The mechanosensory neurons of touch and their mechanisms of activation. Nat Rev Neurosci. 2021;22(9):521–37.

11. Colloca L, Ludman T, Bouhassira D, Baron R, Dickenson AH, Yarnitsky D, et al. Neuropathic pain. Nat Rev Dis Primers. 2017;3:17002.

12. Churko JM, Laird DW. Gap junction remodeling in skin repair following wounding and disease. Physiology (Bethesda). 2013;28(3):190–8.

13. Woo SH, Lumpkin EA, Patapoutian A. Merkel cells and neurons keep in touch. Trends Cell Biol. 2015;25(2):74–81.

14. Shiohara T, Moriya N. Epidermal T cells: their functional role and disease relevance for dermatologists. J Invest Dermatol. 1997;109(3):271–5.

15. Di Meglio P, Perera GK, Nestle FO. The multitasking organ: recent insights into skin immune function. Immunity. 2011;35(6):857–69.

16. Nestle FO, Di Meglio P, Qin JZ, Nickoloff BJ. Skin immune sentinels in health and disease. Nature reviews Immunology. 2009;9(10):679–91.

17. Ho AW, Kupper TS. T cells and the skin: from protective immunity to inflammatory skin disorders. Nature reviews Immunology. 2019;19(8):490–502.

18. Calvo M, Dawes JM, Bennett DL. The role of the immune system in the generation of neuropathic pain. Lancet Neurol. 2012;11(7):629–42.

19. Vincent AM, Callaghan BC, Smith AL, Feldman EL. Diabetic neuropathy: cellular mechanisms as therapeutic targets. Nat Rev Neurol. 2011;7(10):573–83.

20. Uceyler N, Rogausch JP, Toyka KV, Sommer C. Differential expression of cytokines in painful and painless neuropathies. Neurology. 2007;69(1):42–9.

21. Kampoli AM, Tousoulis D, Briasoulis A, Latsios G, Papageorgiou N, Stefanadis C. Potential pathogenic inflammatory mechanisms of endothelial dysfunction induced by type 2 diabetes mellitus. Current pharmaceutical design. 2011;17(37):4147–58.

22. Sjoholm A, Nystrom T. Endothelial inflammation in insulin resistance. Lancet. 2005;365(9459):610–2.

23. Gylfadottir SS, Itani M, Kristensen AG, Tankisi H, Jensen TS, Sindrup SH, et al. Analysis of Macrophages and Peptidergic Fibers in the Skin of Patients With Painful Diabetic Polyneuropathy. Neurol Neuroimmunol Neuroinflamm. 2022;9(1).

24. Langerhans P. Ueber die Nerven der menschlichen Haut. Archiv für pathologische Anatomie und Physiologie und für klinische Medicin. 1868;44(2):325–37.

25. West HC, Bennett CL. Redefining the Role of Langerhans Cells As Immune Regulators within the Skin. Front Immunol. 2017;8:1941.

26. Doebel T, Voisin B, Nagao K. Langerhans Cells - The Macrophage in Dendritic Cell Clothing. Trends Immunol. 2017;38(11):817–28.

27. Kaplan DH. Ontogeny and function of murine epidermal Langerhans cells. Nat Immunol. 2017;18(10):1068–75.

28. Satpathy AT, Wu X, Albring JC, Murphy KM. Re(de)fining the dendritic cell lineage. Nat Immunol. 2012;13(12):1145–54.

29. Bos JD, Kapsenberg ML. The skin immune system Its cellular constituents and their interactions. Immunol Today. 1986;7(7-8):235–40.

30. Cui A, Huang T, Li S, Ma A, Perez JL, Sander C, et al. Dictionary of immune responses to cytokines at single-cell resolution. Nature. 2024;625(7994):377–84.

31. Uceyler N, Vollert J, Broll B, Riediger N, Langjahr M, Saffer N, et al. Sensory profiles and skin innervation of patients with painful and painless neuropathies. Pain. 2018;159(9):1867–76.

32. Hur J, Sullivan KA, Pande M, Hong Y, Sima AA, Jagadish HV, et al. The identification of gene expression profiles associated with progression of human diabetic neuropathy. Brain. 2011;134(Pt 11):3222–35.

33. Miller RJ, Jung H, Bhangoo SK, White FA. Cytokine and chemokine regulation of sensory neuron function. Handb Exp Pharmacol. 2009(194):417–49.

34. Casanova-Molla J, Morales M, Planas-Rigol E, Bosch A, Calvo M, Grau-Junyent JM, et al. Epidermal Langerhans cells in small fiber neuropathies. Pain. 2012;153(5):982–9.

35. Doss AL, Smith PG. Langerhans cells regulate cutaneous innervation density and mechanical sensitivity in mouse footpad. Neurosci Lett. 2014;578:55–60.

36. Dauch JR, Bender DE, Luna-Wong LA, Hsieh W, Yanik BM, Kelly ZA, et al. Neurogenic factor-induced Langerhans cell activation in diabetic mice with mechanical allodynia. J Neuroinflammation. 2013;10:64.

37. Zhang S, Edwards TN, Chaudhri VK, Wu J, Cohen JA, Hirai T, et al. Nonpeptidergic neurons suppress mast cells via glutamate to maintain skin homeostasis. Cell. 2021;184(8):2151–66 e16.

38. George DS, Hackelberg S, Jayaraj ND, Ren D, Edassery SL, Rathwell C, et al. Mitochondrial calcium uniporter deletion prevents painful diabetic neuropathy by restoring mitochondrial morphology and dynamics. Pain. 2021.

39. George DS, Jayaraj ND, Pacifico P, Ren D, Sriram N, Miller RE, et al. The Mas-related G protein-coupled receptor d (Mrgprd) mediates pain hypersensitivity in painful diabetic neuropathy. Pain. 2024.

40. Jayaraj ND, Bhattacharyya BJ, Belmadani AA, Ren D, Rathwell CA, Hackelberg S, et al. Reducing CXCR4-mediated nociceptor hyperexcitability reverses painful diabetic neuropathy. J Clin Invest. 2018;128(6):2205–25.

41. Menichella DM, Abdelhak B, Ren D, Shum A, Frietag C, Miller RJ. CXCR4 chemokine receptor signaling mediates pain in diabetic neuropathy. Molecular pain. 2014;10:42.

42. Obrosova IG, Ilnytska O, Lyzogubov VV, Pavlov IA, Mashtalir N, Nadler JL, et al. High-fat diet induced neuropathy of pre-diabetes and obesity: effects of “healthy” diet and aldose reductase inhibition. Diabetes. 2007;56(10):2598–608.

43. Menichella DM, Jayaraj ND, Wilson HM, Ren D, Flood K, Wang XQ, et al. Ganglioside GM3 synthase depletion reverses neuropathic pain and small fiber neuropathy in diet-induced diabetic mice. Molecular pain. 2016;12.

44. Liu X, Zhu R, Luo Y, Wang S, Zhao Y, Qiu Z, et al. Distinct human Langerhans cell subsets orchestrate reciprocal functions and require different developmental regulation. Immunity. 2021;54(10):2305–20 e11.

45. Ouchi T, Nakato G, Udey MC. EpCAM Expressed by Murine Epidermal Langerhans Cells Modulates Immunization to an Epicutaneously Applied Protein Antigen. J Invest Dermatol. 2016;136(8):1627–35.

46. Ozawa H, Aiba S, Nakagawa S, Tagami H. Murine epidermal Langerhans cells express CD48, which is a counter-receptor for mouse CD2. Arch Dermatol Res. 1995;287(6):524–8.

47. Sere K, Baek JH, Ober-Blobaum J, Muller-Newen G, Tacke F, Yokota Y, et al. Two distinct types of Langerhans cells populate the skin during steady state and inflammation. Immunity. 2012;37(5):905–16.

48. Egenolf N, Zu Altenschildesche CM, Kress L, Eggermann K, Namer B, Gross F, et al. Diagnosing small fiber neuropathy in clinical practice: a deep phenotyping study. Ther Adv Neurol Disord. 2021;14:17562864211004318.

49. Hoeijmakers JG, Faber CG, Lauria G, Merkies IS, Waxman SG. Small-fibre neuropathies--advances in diagnosis, pathophysiology and management. Nat Rev Neurol. 2012;8(7):369–79.

50. Lauria G, Bakkers M, Schmitz C, Lombardi R, Penza P, Devigili G, et al. Intraepidermal nerve fiber density at the distal leg: a worldwide normative reference study. J Peripher Nerv Syst. 2010;15(3):202–7.

51. Merad M, Ginhoux F, Collin M. Origin, homeostasis and function of Langerhans cells and other langerin-expressing dendritic cells. Nature reviews Immunology. 2008;8(12):935–47.

52. Kaplan DH, Jenison MC, Saeland S, Shlomchik WD, Shlomchik MJ. Epidermal langerhans cell-deficient mice develop enhanced contact hypersensitivity. Immunity. 2005;23(6):611–20.

53. Zhang H, Lecker I, Collymore C, Dokova A, Pham MC, Rosen SF, et al. Cage-lid hanging behavior as a translationally relevant measure of pain in mice. Pain. 2021;162(5):1416–25.

54. Subramanian A, Tamayo P, Mootha VK, Mukherjee S, Ebert BL, Gillette MA, et al. Gene set enrichment analysis: a knowledge-based approach for interpreting genome-wide expression profiles. Proc Natl Acad Sci U S A. 2005;102(43):15545–50.

55. Liberzon A, Birger C, Thorvaldsdottir H, Ghandi M, Mesirov JP, Tamayo P. The Molecular Signatures Database (MSigDB) hallmark gene set collection. Cell Syst. 2015;1(6):417–25.

56. Ferrer IR, West HC, Henderson S, Ushakov DS, Santos ESP, Strid J, et al. A wave of monocytes is recruited to replenish the long-term Langerhans cell network after immune injury. Sci Immunol. 2019;4(38).

57. Singh TP, Zhang HH, Borek I, Wolf P, Hedrick MN, Singh SP, et al. Monocyte-derived inflammatory Langerhans cells and dermal dendritic cells mediate psoriasis-like inflammation. Nat Commun. 2016;7:13581.

58. Hovav AH. Mucosal and Skin Langerhans Cells - Nurture Calls. Trends Immunol. 2018;39(10):788–800.

59. Reynolds G, Vegh P, Fletcher J, Poyner EFM, Stephenson E, Goh I, et al. Developmental cell programs are co-opted in inflammatory skin disease. Science. 2021;371(6527).

60. Guilliams M, Mildner A, Yona S. Developmental and Functional Heterogeneity of Monocytes. Immunity. 2018;49(4):595–613.

61. Kubo A, Nagao K, Yokouchi M, Sasaki H, Amagai M. External antigen uptake by Langerhans cells with reorganization of epidermal tight junction barriers. The Journal of experimental medicine. 2009;206(13):2937–46.

62. McArdel SL, Terhorst C, Sharpe AH. Roles of CD48 in regulating immunity and tolerance. Clin Immunol. 2016;164:10–20.

63. Wang J, Li X, Qiang X, Yin X, Guo L. Analyzing the expression and clinical significance of CENPE in gastric cancer. BMC medical genomics. 2024;17(1):119.

64. Silva JR, Iftinca M, Fernandes Gomes FI, Segal JP, Smith OMA, Bannerman CA, et al. Skin-resident dendritic cells mediate postoperative pain via CCR4 on sensory neurons. Proc Natl Acad Sci U S A. 2022;119(4).

65. Ghasemlou N, Chiu IM, Julien JP, Woolf CJ. CD11b+Ly6G-myeloid cells mediate mechanical inflammatory pain hypersensitivity. Proc Natl Acad Sci U S A. 2015;112(49):E6808–17.

66. Kupari J, Ernfors P. Molecular taxonomy of nociceptors and pruriceptors. Pain. 2023.

67. Kupari J, Usoskin D, Parisien M, Lou D, Hu Y, Fatt M, et al. Single cell transcriptomics of primate sensory neurons identifies cell types associated with chronic pain. Nat Commun. 2021;12(1):1510.

68. Yang D, Jacobson A, Meerschaert KA, Sifakis JJ, Wu M, Chen X, et al. Nociceptor neurons direct goblet cells via a CGRP-RAMP1 axis to drive mucus production and gut barrier protection. Cell. 2022;185(22):4190–205 e25.

69. Sun JH, Yang B, Donnelly DF, Ma C, LaMotte RH. MCP-1 enhances excitability of nociceptive neurons in chronically compressed dorsal root ganglia. Journal of neurophysiology. 2006;96(5):2189–99.

70. White FA, Feldman P, Miller RJ. Chemokine signaling and the management of neuropathic pain. Molecular interventions. 2009;9(4):188–95.

71. Jin S, Guerrero-Juarez CF, Zhang L, Chang I, Ramos R, Kuan CH, et al. Inference and analysis of cell-cell communication using CellChat. Nat Commun. 2021;12(1):1088.

72. Deuel TF, Zhang N, Yeh HJ, Silos-Santiago I, Wang ZY. Pleiotrophin: a cytokine with diverse functions and a novel signaling pathway. Arch Biochem Biophys. 2002;397(2):162–71.

73. Pacifico P, Coy-Dibley JS, Miller RJ, Menichella DM. Peripheral mechanisms of peripheral neuropathic pain. Front Mol Neurosci. 2023;16:1252442.

74. Kuhn JA, Vainchtein ID, Braz J, Hamel K, Bernstein M, Craik V, et al. Regulatory T-cells inhibit microglia-induced pain hypersensitivity in female mice. Elife. 2021;10.

75. Mogil JS. Sex differences in pain and pain inhibition: multiple explanations of a controversial phenomenon. Nat Rev Neurosci. 2012;13(12):859–66.

76. Mogil JS. Qualitative sex differences in pain processing: emerging evidence of a biased literature. Nat Rev Neurosci. 2020;21(7):353–65.

77. Klein SL, Flanagan KL. Sex differences in immune responses. Nature reviews Immunology. 2016;16(10):626–38.

78. Scholz J, Woolf CJ. The neuropathic pain triad: neurons, immune cells and glia. Nat Neurosci. 2007;10(11):1361–8.

79. Kashem SW, Haniffa M, Kaplan DH. Antigen-Presenting Cells in the Skin. Annual review of immunology. 2017;35:469–99.

80. Nishibu A, Ward BR, Jester JV, Ploegh HL, Boes M, Takashima A. Behavioral responses of epidermal Langerhans cells in situ to local pathological stimuli. J Invest Dermatol. 2006;126(4):787–96.

81. Stoitzner P, Pfaller K, Stossel H, Romani N. A close-up view of migrating Langerhans cells in the skin. J Invest Dermatol. 2002;118(1):117–25.

82. Kress L, Egenolf N, Sommer C, Uceyler N. Cytokine expression profiles in white blood cells of patients with small fiber neuropathy. BMC Neurosci. 2023;24(1):1.

83. Uceyler N, Schafers M, Sommer C. Mode of action of cytokines on nociceptive neurons. Experimental brain research Experimentelle Hirnforschung Experimentation cerebrale. 2009;196(1):67–78.

84. Costigan M, Scholz J, Woolf CJ. Neuropathic pain: a maladaptive response of the nervous system to damage. Annu Rev Neurosci. 2009;32:1–32.

85. Abbadie C. Chemokines, chemokine receptors and pain. Trends Immunol. 2005;26(10):529–34.

86. Abbadie C, Lindia JA, Cumiskey AM, Peterson LB, Mudgett JS, Bayne EK, et al. Impaired neuropathic pain responses in mice lacking the chemokine receptor CCR2. Proc Natl Acad Sci U S A. 2003;100(13):7947–52.

87. Tanaka T, Minami M, Nakagawa T, Satoh M. Enhanced production of monocyte chemoattractant protein-1 in the dorsal root ganglia in a rat model of neuropathic pain: possible involvement in the development of neuropathic pain. Neurosci Res. 2004;48(4):463–9.

88. White FA, Sun J, Waters SM, Ma C, Ren D, Ripsch M, et al. Excitatory monocyte chemoattractant protein-1 signaling is up-regulated in sensory neurons after chronic compression of the dorsal root ganglion. Proc Natl Acad Sci U S A. 2005;102(39):14092–7.

89. White FA, Jung H, Miller RJ. Chemokines and the pathophysiology of neuropathic pain. Proc Natl Acad Sci U S A. 2007;104(51):20151–8.

90. White FA, Miller RJ. Insights into the regulation of chemokine receptors by molecular signaling pathways: functional roles in neuropathic pain. Brain, behavior, and immunity. 2010;24(6):859–65.

91. Hakim S, Jain A, Petrova V, Indajang J, Kawaguchi R, Wang Q, et al. Macrophages protect against sensory axon degeneration in diabetic neuropathy. bioRxiv. 2024.

92. Griffiths CE, Dearman RJ, Cumberbatch M, Kimber I. Cytokines and Langerhans cell mobilisation in mouse and man. Cytokine. 2005;32(2):67–70.

93. Miyoshi K, Obata K, Kondo T, Okamura H, Noguchi K. Interleukin-18-mediated microglia/astrocyte interaction in the spinal cord enhances neuropathic pain processing after nerve injury. The Journal of neuroscience : the official journal of the Society for Neuroscience. 2008;28(48):12775–87.

94. Pilat D, Piotrowska A, Rojewska E, Jurga A, Slusarczyk J, Makuch W, et al. Blockade of IL-18 signaling diminished neuropathic pain and enhanced the efficacy of morphine and buprenorphine. Molecular and cellular neurosciences. 2016;71:114–24.

95. Moriwaki Y, Yamamoto T, Shibutani Y, Aoki E, Tsutsumi Z, Takahashi S, et al. Elevated levels of interleukin-18 and tumor necrosis factor-alpha in serum of patients with type 2 diabetes mellitus: relationship with diabetic nephropathy. Metabolism. 2003;52(5):605–8.

96. Gateva A, Kamenov Z, Karamfilova V, Assyov Y, Velikova T, El-Darawish Y, et al. Higher levels of IL-18 in patients with prediabetes compared to obese normoglycaemic controls. Arch Physiol Biochem. 2020;126(5):449–52.

97. Ju J, Li Z, Jia X, Peng X, Wang J, Gao F. Interleukin-18 in chronic pain: Focus on pathogenic mechanisms and potential therapeutic targets. Pharmacol Res. 2024;201:107089.

98. Bautista J, Chandrasekhar A, Komirishetty PK, Duraikannu A, Zochodne DW. Regenerative plasticity of intact human skin axons. Journal of the neurological sciences. 2020;417:117058.

99. Gangadharan V, Zheng H, Taberner FJ, Landry J, Nees TA, Pistolic J, et al. Neuropathic pain caused by miswiring and abnormal end organ targeting. Nature. 2022;606(7912):137–45.

100. Pinho-Ribeiro FA, Chiu IM. Nociceptor nerves set the stage for skin immunity. Cell Res. 2019;29(11):877–8.

101. Kruger RP, Aurandt J, Guan KL. Semaphorins command cells to move. Nat Rev Mol Cell Biol. 2005;6(10):789–800.

102. Tamagnone L, Artigiani S, Chen H, He Z, Ming GI, Song H, et al. Plexins are a large family of receptors for transmembrane, secreted, and GPI-anchored semaphorins in vertebrates. Cell. 1999;99(1):71–80.

103. Pasterkamp RJ. Getting neural circuits into shape with semaphorins. Nat Rev Neurosci. 2012;13(9):605–18.

104. Suzuki K, Kumanogoh A, Kikutani H. Semaphorins and their receptors in immune cell interactions. Nat Immunol. 2008;9(1):17–23.

105. Moretti S, Procopio A, Boemi M, Catalano A. Neuronal semaphorins regulate a primary immune response. Curr Neurovasc Res. 2006;3(4):295–305.

106. Kang S, Nakanishi Y, Kioi Y, Okuzaki D, Kimura T, Takamatsu H, et al. Semaphorin 6D reverse signaling controls macrophage lipid metabolism and anti-inflammatory polarization. Nat Immunol. 2018;19(6):561–70.

107. Witherden DA, Watanabe M, Garijo O, Rieder SE, Sarkisyan G, Cronin SJ, et al. The CD100 receptor interacts with its plexin B2 ligand to regulate epidermal gammadelta T cell function. Immunity. 2012;37(2):314–25.

108. Paldy E, Simonetti M, Worzfeld T, Bali KK, Vicuna L, Offermanns S, et al. Semaphorin 4C Plexin-B2 signaling in peripheral sensory neurons is pronociceptive in a model of inflammatory pain. Nat Commun. 2017;8(1):176.

109. Takamatsu H, Okuno T, Kumanogoh A. Regulation of immune cell responses by semaphorins and their receptors. Cell Mol Immunol. 2010;7(2):83–8.

110. Takamatsu H, Takegahara N, Nakagawa Y, Tomura M, Taniguchi M, Friedel RH, et al. Semaphorins guide the entry of dendritic cells into the lymphatics by activating myosin II. Nat Immunol. 2010;11(7):594–600.

111. Zhou X, Wahane S, Friedl MS, Kluge M, Friedel CC, Avrampou K, et al. Microglia and macrophages promote corralling, wound compaction and recovery after spinal cord injury via Plexin-B2. Nat Neurosci. 2020;23(3):337–50.

112. Tansley S, Uttam S, Urena Guzman A, Yaqubi M, Pacis A, Parisien M, et al. Single-cell RNA sequencing reveals time- and sex-specific responses of mouse spinal cord microglia to peripheral nerve injury and links ApoE to chronic pain. Nat Commun. 2022;13(1):843.

113. Dorfman MD, Krull JE, Douglass JD, Fasnacht R, Lara-Lince F, Meek TH, et al. Sex differences in microglial CX3CR1 signalling determine obesity susceptibility in mice. Nat Commun. 2017;8:14556.

114. Black JA, Waxman SG. Noncanonical roles of voltage-gated sodium channels. Neuron. 2013;80(2):280–91.

115. Eder C. Ion channels in monocytes and microglia/brain macrophages: promising therapeutic targets for neurological diseases. Journal of neuroimmunology. 2010;224(1-2):51–5.

116. Kis-Toth K, Hajdu P, Bacskai I, Szilagyi O, Papp F, Szanto A, et al. Voltage-gated sodium channel Nav1.7 maintains the membrane potential and regulates the activation and chemokine-induced migration of a monocyte-derived dendritic cell subset. Journal of immunology. 2011;187(3):1273–80.

117. Truini A, Cruccu G. How diagnostic tests help to disentangle the mechanisms underlying neuropathic pain symptoms in painful neuropathies. Pain. 2016;157 Suppl 1:S53–S9.

118. Devigili G, Rinaldo S, Lombardi R, Cazzato D, Marchi M, Salvi E, et al. Diagnostic criteria for small fibre neuropathy in clinical practice and research. Brain. 2019;142(12):3728–36.

