## Supplementary material for "Epidermal Langerhans Cells Drive Painful Diabetic Neuropathy in a Sex-dependent Manner": Suppl. Materials includes extended methods sections and supplementary figures

**SUPPLEMENTARY MATERIALS**

### METHODS

#### Ethical statements

This research complies with all relevant ethical regulations.

All animal care protocols and experiments were reviewed and approved by the Institutional Animal Care and Use Committee (IACUC) of Northwestern University.

Human tissues were collected from healthy volunteers and patients and the study was approved by the local Ethical Committee of the Fondazione IRCCS Istituto Neurologico 'Carlo Besta' of Milan (FINCB), Italy.

#### Animals

Animals were housed on a 12-hour light/12-hour dark cycle with ad libitum access to food and water. Adult wild-type male and female mice between 6 and 8 weeks were adopted for most of the experiments. Both male and female were fed a high-fat diet (HFD 42% fat - EnvigoTD88137, Envigo, Madison, WI) or a regular diet (RD 11% fat) for 10 weeks and then a glucose tolerance test was performed as first described in Menichella et al. 2016(1). After fasting, mice were injected with a 45% D-glucose solution (2mg glucose/g body weight) and blood glucose was measured at 30, 60, and 120 minutes after injection. To compare "diabetic" versus "non-diabetic" HFD mice, we set the cutoff for diabetes ≥140 mg/dl for male mice(2) and ≥130 mg/dl for female mice at 2 SD above the mean for glucose 120 minutes after glucose challenge. In addition to wild-type mice, we used the humanized Langerin-diphtheria toxin receptor mouse line(3) (hLangerin-DTR), kindly provided by D. Kaplan at Pittsburg University, and the reporter Nav1.8-dtTomato mouse line, previously described in Jayaraj et al., 2018(2).

**Skin biopsies**

All subjects underwent clinical examination, skin biopsy and gave informed consent to participate in the study. For the present study, 15 patients with diabetic neuropathy (7 females and 8 males) and 9 healthy controls (4 females and 5 males) were enrolled. Clinical features were collected using an established protocol previously described(4). Pain intensity was measured as the average score of the last 3 weeks using the pain intensity numerical rating scale (PI-NRS).

**Single-cell dissociation from paw epidermis**

Glabrous hind paw skin was harvested from the mouse and incubated in Dispase (2.3 mg/ml, Sigma-Aldrich D4693) overnight at 4C. The following day, the epidermis was gently peeled off from the dermis and incubated in TrypLE Express (Gibco 12604-013) for 10 minutes at 37C. Single cells were dissociated using gentle agitation, filtered with a 30 µm cell strainer into a sterile tube and then resuspended in 154CF epidermal medium (Gibco M154CF500) or in RPMI 1640 (Gibco 11875093)

**Cell sorting and primary culture of epidermal LCs.**

Single cells isolated from the paw epidermis and collected in RPMI 1640 were centrifuged at 300g for 5 minutes at 4C. The cell pellet was resuspended in FACS buffer (1% BSA in 1xPBS). LCs were selectively labeled using PE anti-human CD207 1:100 (Biolegend,

352203) and APC andi-mouse CD45 1:100 (Biolegend, 103111) for 2hours at 4C in the dark. After staining, the single cells suspension was washed three times with 1xPBS and resuspended in fresh 1XPBS. Given the fragile nature of LCs, the MACSQuant® Tyto® Cell Sorter was used to sort and collect double-positive cells. Aseptic cartridges were employed to load the cells, and only double positive cells CD207+/CD45+ were collected in the middle compartment of the cartridge and subsequently recovered.

The sorted Cd207+/Cd45+ LCs were resuspended in RPMI 1640 enriched with L-glutamine and 2-Mercaptoethanol (2-ME) and plated onto IgG-coated surfaces(5). The plated cells were then stimulated with recombinant mouse TNF-$\alpha$ (R&D, 410-MT) at a final concentration of 125U/mL and recombinant mouse GM-CSF (R&D, 415-ML) at a final concentration of 500U/mL. 50 µL of medium from both stimulated and un-stimulated LCs derived from RD and HFD mice was collected at various time point and stored at -80C for subsequent analysis.

**Immunohistochemistry**

**Whole-mount staining**. Glabrous hind paw skin was harvested from the mouse and incubated in Dispase (2.3 mg/ml, Sigma-Aldrich D4693) overnight at 4C. The following day, the epidermis was gently separated from the dermis. The isolated paw epidermis was fixed with 4% PFA for 30 minutes at room temperature. After fixation, tissues were washed thoroughly with 1XPBS, and the whole tissue was divided in smaller pieces. Only the central area of the hind paw including footpads was processed for histological analysis. The tissue was first permeabilized with 0.2% Triton X-100 in PBS for 15 minutes, and then incubated in a blocking solution containing 10% NDS - 0.1% BSA in 0.1% Triton

X-100-PBS for 1hour at room temperature. The tissue was then incubated with APC anti-mouse/human Cd207 1:100 (Biolegend, 4C7) overnight at 4C and images were acquired using a confocal fluorescence microscope.

**Skin sections**. Glabrous hind paw skin was harvested from the mouse, fixed with 4% PFA for 90 minutes at room temperature, and then stored in 30% sucrose overnight at 4C. The following day, tissue was included in O.C.T. and stored at -80C. 20um cryosections, placed on superfrost plus slides (Thermo Fisher Scientific), were stained with anti-K14 1:500 (906004, Biolegend), anti-K10 1:500 (Ep1607ihcy, Abcam) and anti-CD207 1:100 (4C7, Biolegend). Secondary antibodies Alexa Fluor 488 goat anti-rabbit antibody (Invitrogen, Thermo Fisher Scientific, 1:250), was used.

**Human skin sections.** Skin biopsy samples were taken from the distal leg (10 cm above the lateral malleolus) following the standard procedure(6). Specimens were then fixed in 2% PLP, cryoprotected and cut into 50 um vertical sections. Samples were washed twice in PBS, then kept for an hour in blocking solution (1%BSA + 2% NGS + 0.5% Triton X-100 in PBS). Incubation with mouse anti-CD207 conjugated to Alexa 633 (Biolegend, 1:100) and rabbit anti-PGP9.5 (Proteintech, 1:1500) was performed overnight on room temperature. After two washes in PBS, secondary antibody incubation Goat-Anti-Rabbit 488 (1:2000) was carried out for 1h on RT. Samples were examined with a confocal laser scan microscope (TCS SP8 AOBS; Leica Microsystems) equipped with Ar/Ar-Kr 488 and 633 nm diode lasers. 3 random images were acquired for each sample by using a HC PL APO CS 40x/1.1 Water immersion objective and HyD detectors. Laser intensity and photo multiplier gain for each channel was adjusted to minimize background noise and saturated pixels, and once defined for control conditions, parameters were kept constants for all acquisitions.

**RNAscope in situ hybridization**

RNAscope in situ hybridization multiplex V2 was performed according to the manufacturer's instructions, Advanced Cell Diagnostics (ACD). Twelve-micrometer DRG cryosections from RD and HFD male mice were incubated with probes Mrgprd (417921-C2/C3), Sema4d (498381-C3), Sema6d (565871-C1). More details reported in George et al., 2024(7).

Analysis. DRG sections were analyzed by imaging the whole DRG using Olympus FV10i confocal microscope, and the images were processed using Fiji. Target mRNA expression was measured as average intensity of target dots per cell(7). First the average background intensity (ABI) based on the integrated intensity of a background region in a selected area (ABI5IntDen background/ area of selected background) was calculated. Then, 10 dots per cell (3 cells per section) were considered and the area and integrated intensity of each dot was measured. Lastly, the average intensity per single dot (AISD) was calculated using the formula: AISD = IntDen selected dots - ABI x area of selected dots / tot number of dots. Mean values of the counts from blinded reviewers were plotted and graphed.

**Morphometric analysis**

Images were processed with FIJI and Arivis Pro (rel 4.2). By using the Neurite Tracer pipeline of Arivis Pro we determined the total number of branching points and the volume occupied by LCs. The parameters used in our set up are:

- Method: Probabilistic Reconstructor
- Branch diameter: 0.284-5 um
- Tubularity sensitivity: 12
- Seeding-Tubularity local threshold: 0.2/ Seed Filter: 40%
- Min Terminal section length: 2 mm

**Behavioral test**

*Mechanical allodynia test (von Frey)* To assess mechanical allodynia, the von Frey test was performed as follow. RD and HFD male and female mice were placed on a metal mesh floor under a transparent plastic dome. Following 60-min of habituation, seven different filaments, each with a specific bending force (10, 20, 40, 60, 80, 100, and 120 mN) were applied in ascending order to the plantar surface of the hind paw. Each filament was applied six times with an interstimulus interval between 10-15 seconds. The von Frey withdrawal threshold was defined as the minimum force eliciting a detectable withdrawal response in at least 50% of trials.

*Spontaneous pain test (cage-lid hanging)* To test spontaneous pain, the cage-lid hanging test was adapted from Zhang et al., 2021(8). Briefly, mice were acclimated to the behavioral testing room for 30 minutes before the test. An empty polypropylene cage (dimension: 290x220x140mm) with a slightly modified lid was used for the test. Each mouse was placed on the underside of the inverted lid, and the time to fall was recorded.

The test was repeated three times for each mouse with a maximum cutoff time of 30 seconds.

Researchers conducting behavioral tests and endpoint analyses were blinded to the experimental conditions.

**scRNAseq**

*Sample processing and raw data analysis*

For the scRNAseq of paw epidermis of three adult RD and HFD male and female mice, we generated a single-cell suspensions as described in "*Single-cell dissociation from paw epidermis*" paragraph. We assessed cell viability using an automated cell counter and for all samples we achieved >90% cell viability.

For the sequencing, we choose the droplet-based method by 10X Genomics where each droplet should contain a single cell. Libraries were prepared by NUSeq Core at Northwestern University according to 10X Genomics instructions. Sequencing was performed on an Illumina platform, and FASTQ raw sequencing data were aligned to mouse transcriptome reference using Cell Ranger toolkit from 10X Genomics. The downstream data analysis and visualization was performed using Seurat 5 pipeline in R.

*Quality control, clustering, and differential gene expression analysis* The standard pre-processing workflow for scRNAseq data in Seurat 5 (version 5.0.0, https://satijalab.org/seurat/)(9) was performed following QC metrics. To remove low-quality cells, potential doublets, or dying cells, filter parameters were applied to both male and female datasets. For paw epidermis of male sample datasets, cells with <200 and >6000 genes, and >5% mitochondrial genes detected were excluded. For paw epidermis of female sample datasets, cells with <500 and >6000 genes, and >5%

mitochondrial genes detected were removed. Seurat 5 in R was used to separately analyze male and female datasets. Features expressions were normalized using Log-normalize and high variable features were identified by FindVariableFeatures function of Seurat package. Principal components analysis was performed using RunPCA, and the top 10 principal components, identified by the cumulative variance analysis, were used for clustering. Uniform Manifold Approximation and Projection (UMAP) was run for clusters' visualization. Differentially expressed genes between RD and HFD groups were identified using a Wilcoxon Rank Sum test, corrected for false discovery rate (FDR), and visualized using the EnhancedVolcano R package. Only genes with log2fold change above 0.25 were considered. The analysis to identify pathways and gene set enrichment was performed using Gene Set Enrichment Analysis (GSEA). Data were visualized using R packages: Seurat, ggplot2, dittoSeq.

*Cell-cell interaction data analysis* To predict intercellular communication, CellChat (v1), a package developed in R(10), was applied to male scRNAseq dataset. CellChat is designed for inference, analysis, and visualization of cell-cell communication from scRNAseq data. The CellChat object created from the Seurat object was aligned to mouse ligand-receptor database to create annotations. The ligand-receptor CellChatDB is based on KEGG (Kyoto Encyclopedia of Genes and Genomes). "computeCommunProb" function was adopted to infer cell-cell communications within the dataset and then standard workflow for visualization was applied to generate graphs.

**Multiplex cytokine analysis**

The cytokine profile of LCs was assessed using the Codeplex Secretome platform by Isoxplexis (Bruker, Cellular Analysis) designed to the molecules of mouse innate immune response. Medium collected from primary LCs culture, as described in the *“Cell sorting and primary culture of epidermal LCs”,* was loaded onto Isoplexis chip (CODEPLEX-2L12-1) for secretome analysis. Cytokines levels were detected and represented as as Relative Fluorescence Unit (RFU) and reported either as Log-Transformed values or pg/mL. The analysis was conducted with the technical support of Bruker specialists.

**Statistics**

All statistical analysis was performed using R studio (2023.12.0+369) or GraphPad Prism (10.0.3). For comparisons between two groups, a two-tailed student’s t-test was applied, and where applicable, adjustments for multiple testing were made and reported in figure legends. For comparisons involving more than two groups, one-way or two-way Analysis of Variance (ANOVA) was performed, followed by post-hoc multiple comparison testing, as reported in figures legends. Pearson correlation analysis was performed to assess the correlation between variables and the correlation coefficient (*r*) and p value were reported within the graphs. Quantification of LCs density, RNAscope and behavioral tests were performed in double-blind. All values are expressed as the mean ± SEM, and a *P* value of less than 0.05 was considered statistically significant.

### DATA AVAILABILITY

All data presented in this study are included in the main text and in supplementary figures. scRNAseq data generated from the paw epidermis of male and female mice will be deposited upon acceptance.


### ACKNOWLEDGEMENTS

The authors thank Dr. Daniel Kaplan for kindly providing h-langerin-DTR mice, and Dr. Rajeshwar Awatramani for insightful discussions. The authors want to acknowledge Dr. Cheryl L. Stucky and Dr. Daniela Salvemini for their thoughtful feedback. This research was partially supported by the Northwestern University Core Centers and funded by the Italian Ministry of Health (RRC). The authors also acknowledge the NUSeq Core for their assistance with the single-cell RNA sequencing, the Pathology Core for Hematoxylin and Eosin staining, the Flow Cytometry Core Facility of the Robert H. Lurie Comprehensive Cancer Center for the assistance with cell sorting. The authors also thank the Immunotherapy Assessment Core at NU, the Flow Cytometry Core at UIC and Bruker scientists for their support with the IsoPlexis experiment. Finally, authors thank the Data Science, Statistics, and Visualization team at Northwestern for their support with data analysis.

This work was supported by NIH R01 NS104295-01 and NIH HEAL initiative supplement R01 NS104295-01 (D.M.M.), NIH R01 AR077691-01 (D.M.M., R.J.M.)

**SUPPLEMENTARY FIGURES**

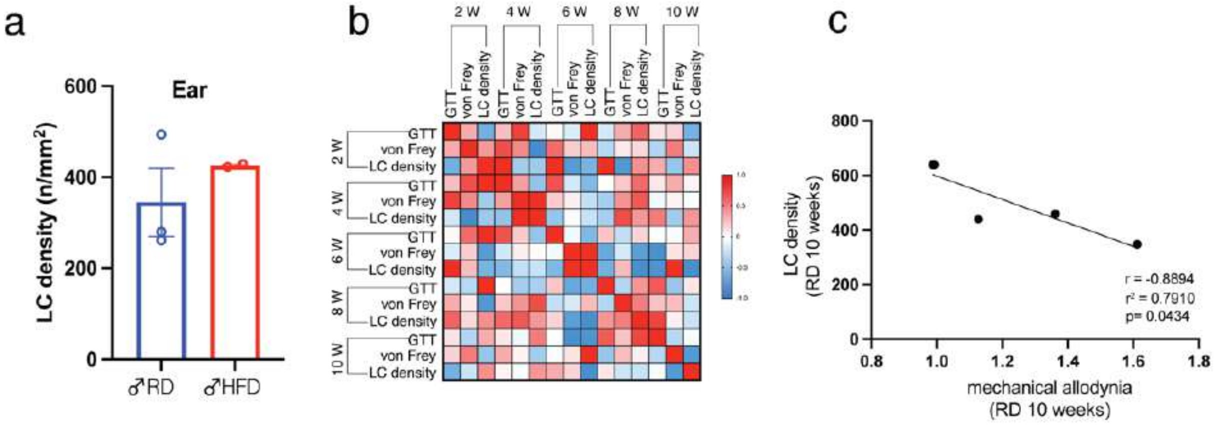


**Suppl. Fig. 1. LCs density in RD mice (a)** LCs density measured as numb of cells/area ($mm^2$) in the ear of male mice. No differences between RD and HFD male animals. Unpaired two tailed t-test with Welch's correction p=0.3940. RD n=3 and HFD n=2. **(b)** Correlation matrix of RD features (GTT, von Frey and LCs) at different time points. **(c)** Negative correlation between LCs density and von Frey in RD male mice. Pearson r coefficient r=-0.8894, $r^2$=0.7910, p= 0.0434(*).

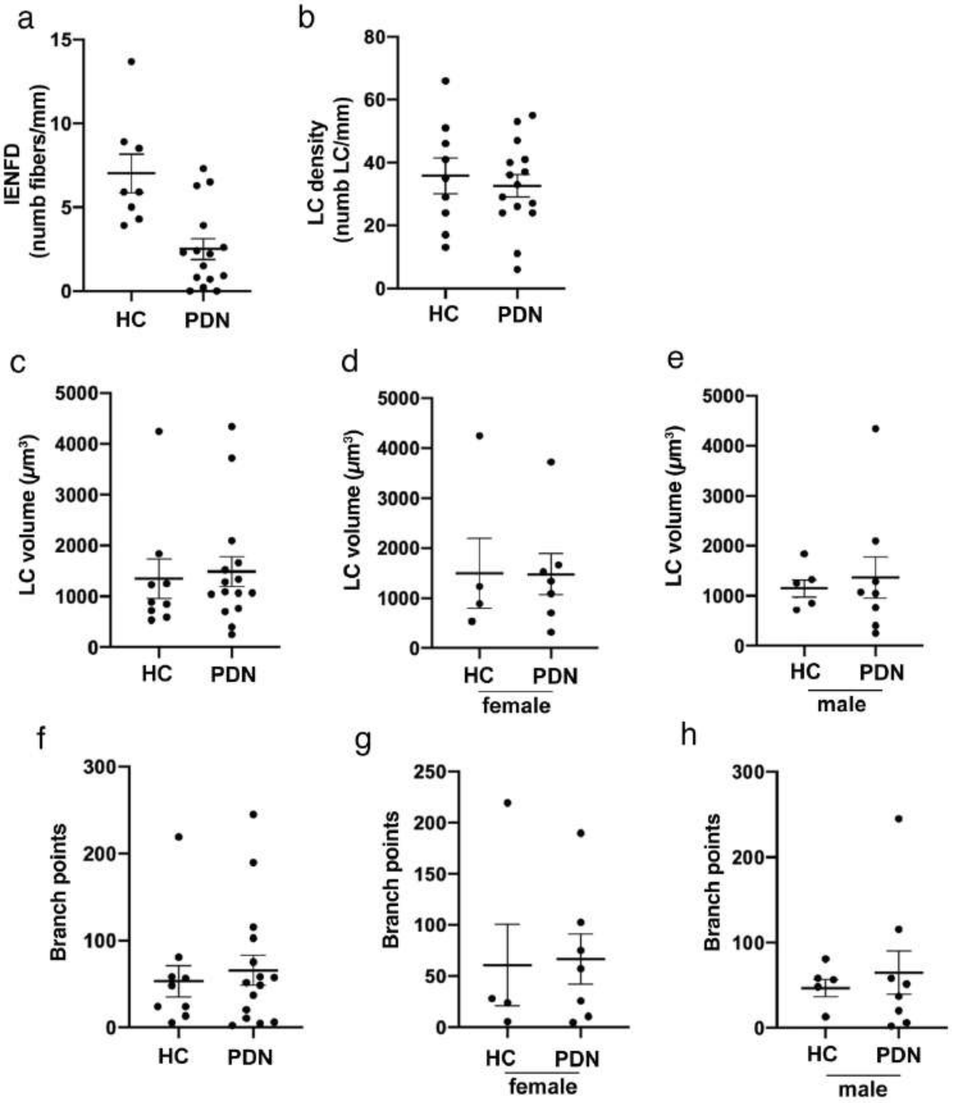


**Suppl. Fig. 2**. **LCs features analysis in PDN patients (a-b)** Plots of the intraepidermal nerve fiber density (IENFD) and the LCs density in the entire cohort of healthy subjects (HC) or diabetic patients (PDN). **(c-e)** LCs volume expressed in $\mu m^3$ in the (c) entire cohort, (d) in female or (e) in male subjects. **(f-h)** branch points of LCs in in the (f) entire cohort, (g) in female or (h) in male subjects. No differences between HC and PDN. Middle lines of the plots represent the mean value, while whiskers are SEM

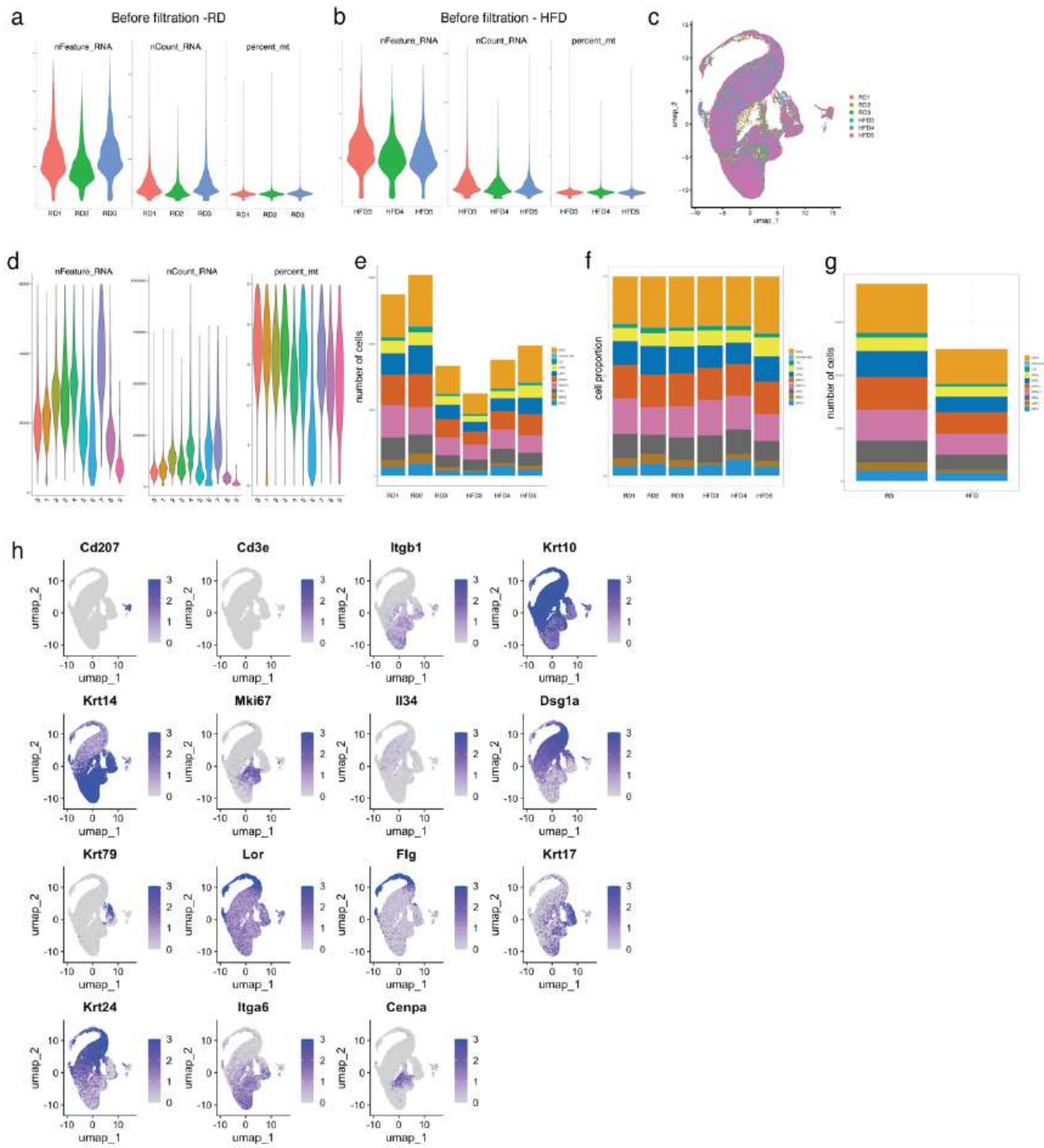


**Suppl. Fig. 3. QC analysis of scRNAseq of RD and HFD paw epidermis (a-b)** QC analysis of RD and HFD samples before applying filtration. **(c)** UMAP dimensionality reduction of each sample from RD and HFD show a full overlap**(d)** Cluster QC analysis after filtration for combined RD and HFD. **(e)** Number of cells in each cluster in each sample: RD1=14781, RD2=16221, RD3=9498, HFD3=7521, HFD4=10306, HFD5=11123 cells. **(f)** Cell proportion in each cluster in each sample. **(g)** Total number of cells in combined samples for RD and HFD: RD=40500 and HFD=28950 cells. **(h)** Feature plots of selected marker genes for cluster identification

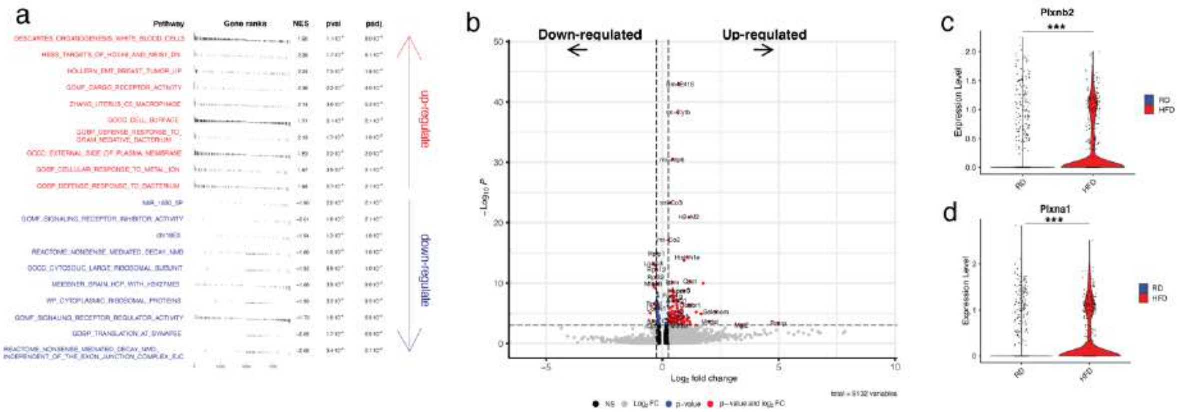


**Suppl. Fig. 4. DEGs in LCs in HFD male mice (a)** GSEA table shows top10 upregulated and the top10 downregulated pathways. GSEA based on false discovery rate (FDR) of 0.25. **(b)** Volcano plot of DEGs identified in Langerhans cells cluster in scRNAseq data from mouse paw epidermis. DEGs adjusted for BH correction. Volcano plot generated with EnhancedVolcano package in R. **(c-d)** Violin Plots show the comparative expression of Plxnb2 (*** p=0.00014) (c) and Plxna1 (*** p=0.00037) (d) in LCs subclusters between RD and HFD. Wilcoxon Rank-Sum test is applied for significance.

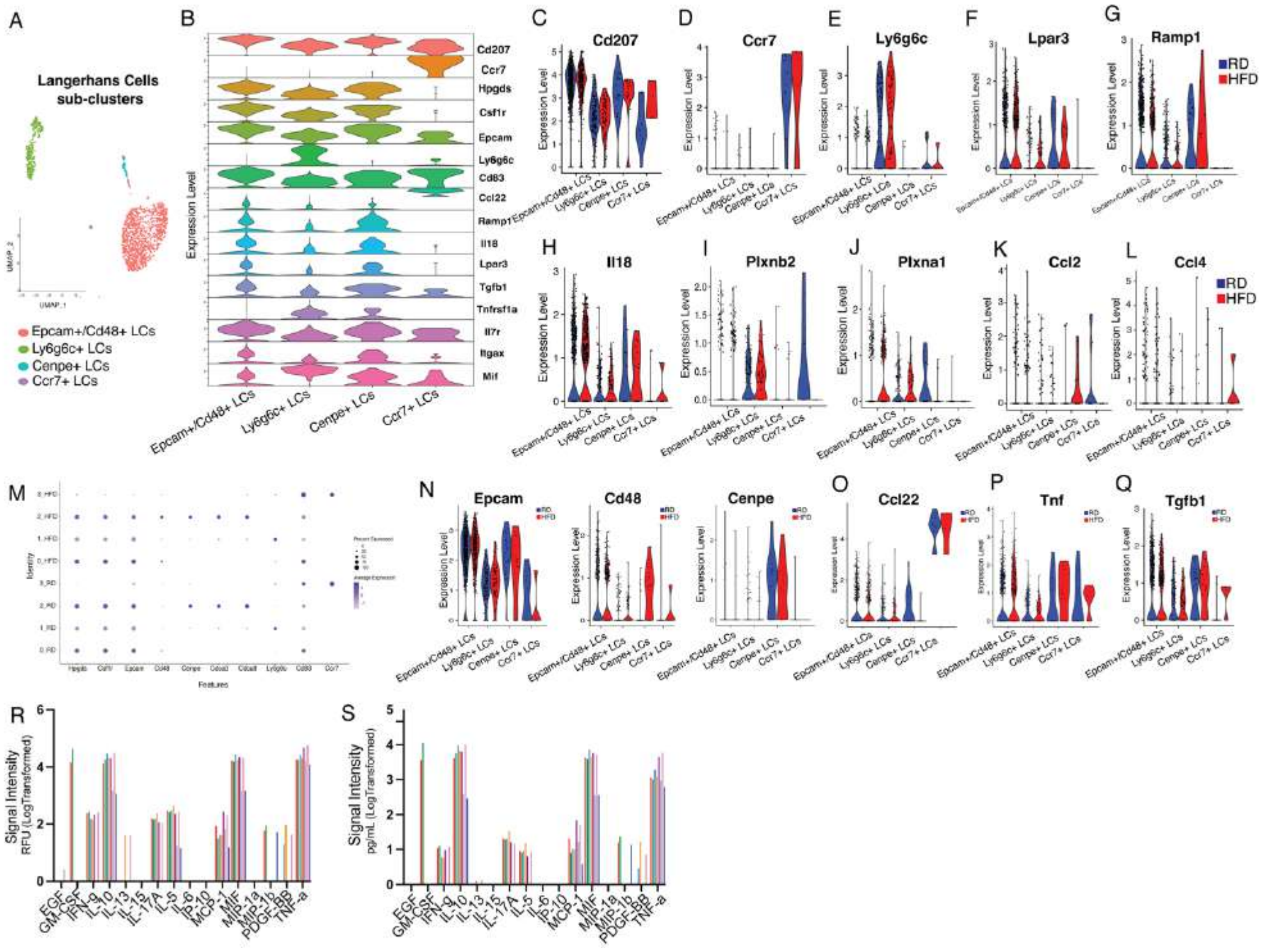


**Suppl. Fig. 5 (A)** UMAP plots of the LCs subclustering showing four distinct groups. **(B)** Stacked violin plot of marker genes adopted to identify LCs subclusters. **(C-H)** ViolinPlot showing the expression of Cd207 (c), Ccr7 (d), Ly6g6c (e), Lpar3 (f), Ramp1 (g) and Il-18 (h) in LCs subclusters. **(I-L)** ViolinPlots show the expression and distribution of plexins molecules Plxnb2 (i) and Plxna1 (j) and chemokine molecules (k) Mcp1/Ccl2 and (l) Mip-1b/Ccl4 in LCs subclusters **(M)** DotPlot shows the expression of genes adopted for subcluster identification **(N-Q)** Violin plots show the differential expression of genes of interested in LCs subclusters. RD=blue, HFD=red. (**R-S)** the high-multiplexing quantitative inflammatory molecule proteomic (CodePlex by Bruker) shows a panel of 16 molecules belonging to the mouse innate immune response and detected in LCs RD and HFD secretome. LCs were stimulated with (1) GM-CSF and the secretome was collected after 12h, (2) TNF-$\alpha$ and the secretome was collected after 12h and 24h and (3) no stimulated and secretome collected after 2h. Molecules are shown as Signal intensity – relative fluorescence unit (RFU and pg/mL, both LogTransformed.

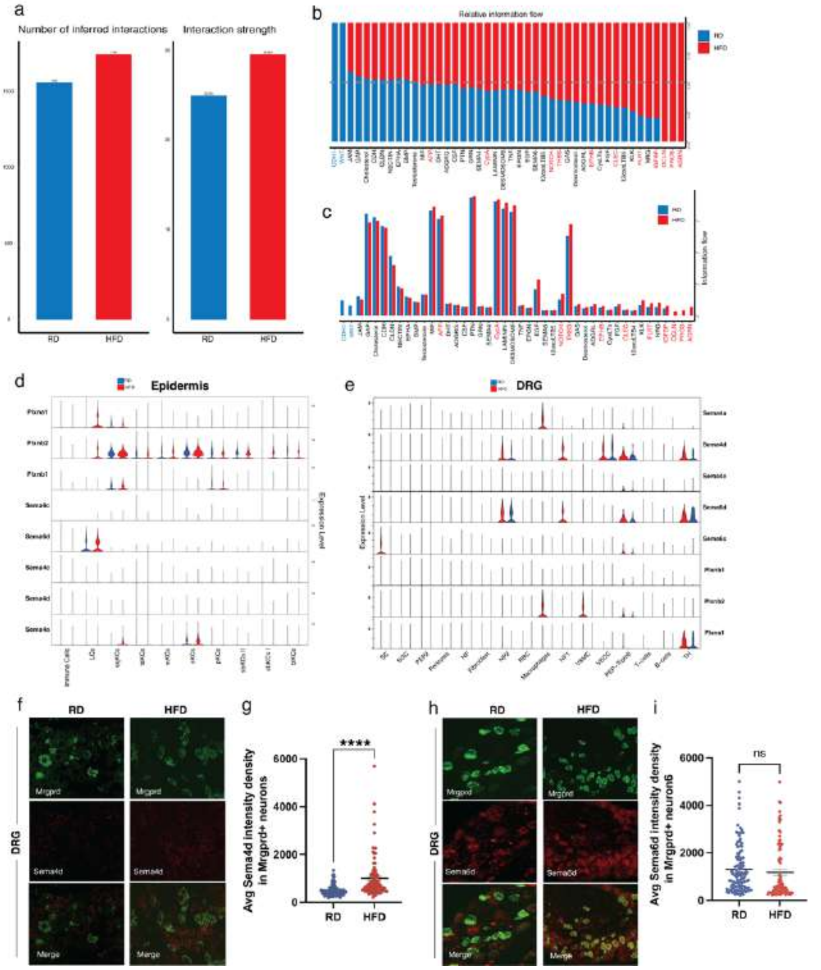


**Suppl. Fig. 6 Sema-Plxn signaling pathways in RD and HFD male mice (a)** CellChat analysis. Bar plots show the numb of inferred interactions and the strength of interactions that are higher in HFD**. (b-c)** Identification of signaling pathways by comparing the information flow of each signaling pathway in each diet condition. **(d)** Stacked VlnPlot shows the expression of Sema4 and Sema6 family members and Plxnb2 and Plxna1 family members in RD and HFD epidermis scRNAseq data. **(e)** Stacked VlnPlot shows the expression of Sema4 and Sema6 family members and Plxnb2 and Plxna1 family members in RD and HFD DRG neurons scRNAseq data. **(f)** Representative images of *in situ* RNAscope of Mrgprd (green) and Sema4d (red) expression in DRG sensory neurons from RD and HFD. **(g)** Quantification of Average Intensity for each Sema4d dot in Mrgprd+ neurons. 30 dots in Mrgprd+ neurons per animals were quantified. Unpaired two-tailed t-Test $p<0.0001$ (****). N=3 animals for each diet condition. **(h)** Representative images of *in situ* RNAscope of Mrgprd (green) and Sema6d (red) expression in DRG sensory neurons from RD and HFD. **(i)** Quantification of Average Intensity for each Sema6d dot in Mrgprd+ neurons. 30 dots in Mrgprd+ neurons per animals were quantified. Unpaired two-tailed t-Test $p=0.4277$ (ns). RD n=4; HFD n=3.

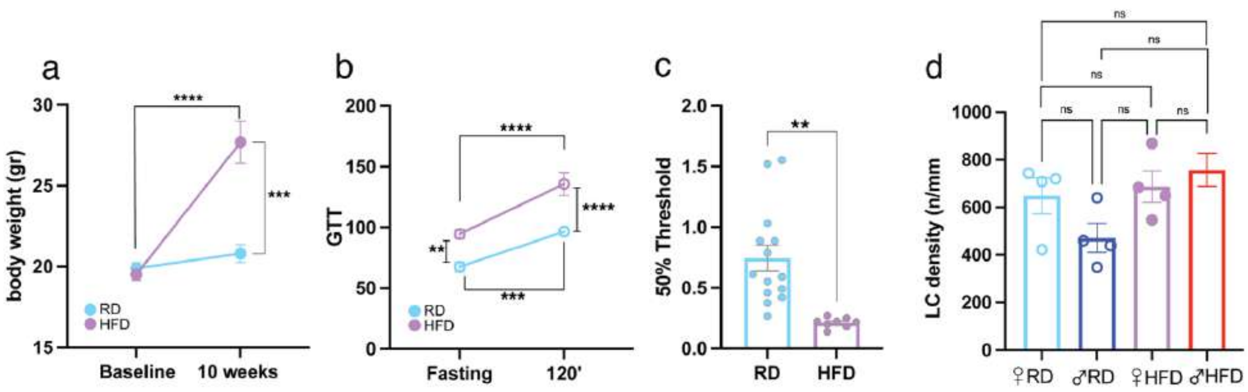


**Suppl. Fig. 7. Phenotypic characterization of HFD in female mice (a)** Body weight (gr) in female mice. n=10 animals per each diet conditions. Two-way ANOVA for multiple comparison. Between RD and HFD: 10 weeks p=0.0002 (***). Within HFD: baseline vs 10weeks p<0.0001. **(b)** GTT in female mice. RD n=14 and HFD n=12. Two-way ANOVA for multiple comparison. Between RD and HFD: Fasting p=0.0011 (**); 120' p<0.0001 (****). Within RD: Fasting vs 120' p=0.0003 (***). Within HFD Fasting vs 120' p<0.0001(****). **(c)** von Frey test. RD n=14 and HFD n=8. Unpaired two-tailed t-Test p=0.0014 (**). **(d)** LCs' density compared among all groups. One-way ANOVA for multiple comparisons. RD female vs. RD male p=ns; HFD female vs. HFD male p=ns; RD female vs. HFD female p=ns; RD female vs. HFD male p=ns; RD male vs. HFD male p=0.045 (*); RD male vs HFD female p=ns.

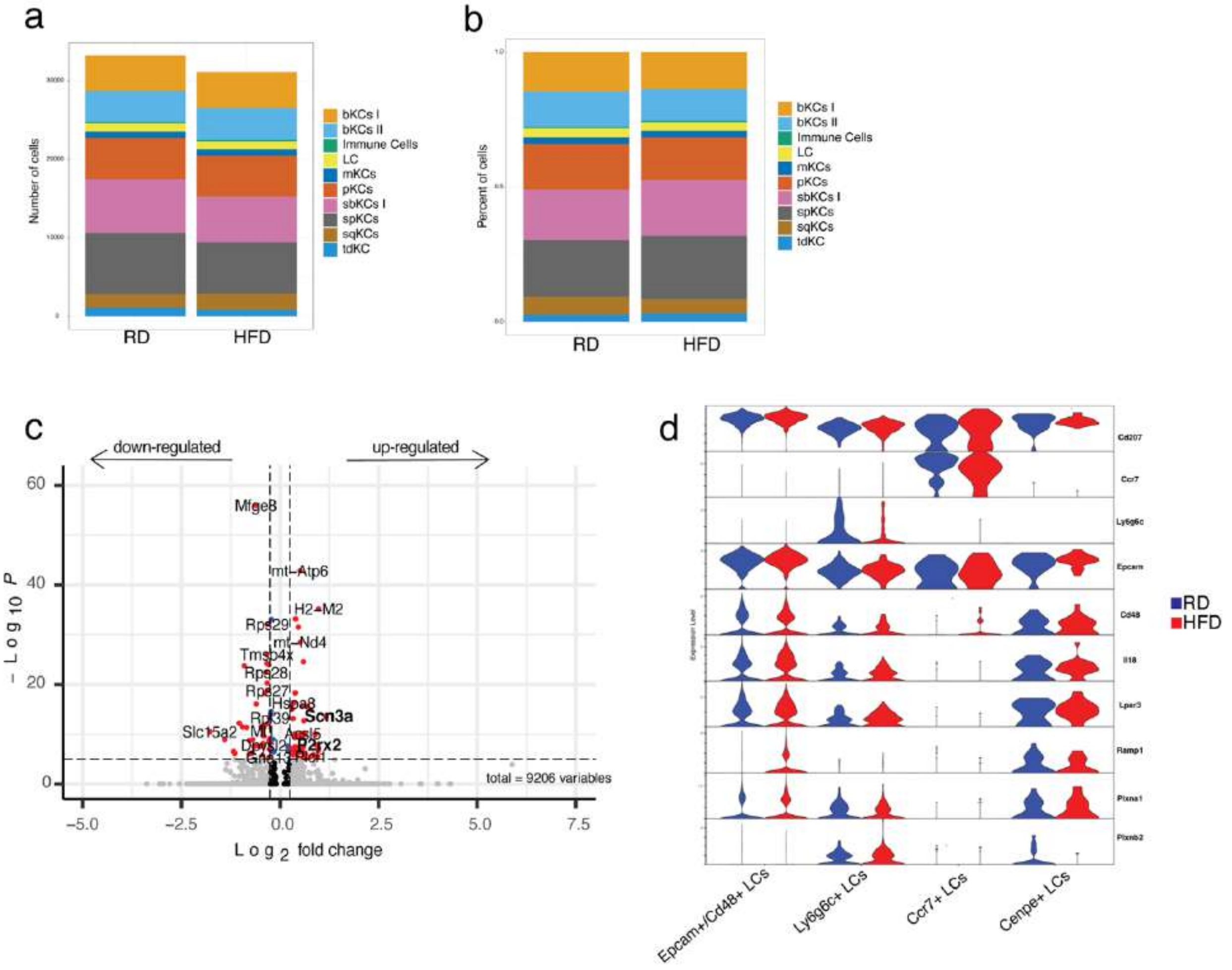


**Suppl. Fig. 8. QC analysis of scRNAseq of paw epidermis of RD and HFD female mice (a-b)** QC analysis measuring the number of cells and the percentage of cells in each cluster of scRNAseq paw epidermis of RD and HFD female mice. **(c)** Volcano plot of DEGs identified in LCs cluster in scRNAseq data from female mouse paw epidermis. DEGs adjusted for BH correction. Volcano plot generated with EnhancedVolcano package in R. **(d)** Stacked violin plots show the expression genes adopted to identified LCs subclusters and found different in male LCs subcluster.

1. Menichella DM, Jayaraj ND, Wilson HM, Ren D, Flood K, Wang XQ, et al. Ganglioside GM3 synthase depletion reverses neuropathic pain and small fiber neuropathy in diet-induced diabetic mice. Molecular pain. 2016;12.

2. Jayaraj ND, Bhattacharyya BJ, Belmadani AA, Ren D, Rathwell CA, Hackelberg S, et al. Reducing CXCR4-mediated nociceptor hyperexcitability reverses painful diabetic neuropathy. J Clin Invest. 2018;128(6):2205-25.

3. Zhang S, Edwards TN, Chaudhri VK, Wu J, Cohen JA, Hirai T, et al. Nonpeptidergic neurons suppress mast cells via glutamate to maintain skin homeostasis. Cell. 2021;184(8):2151-66 e16.

4. Devigili G, Rinaldo S, Lombardi R, Cazzato D, Marchi M, Salvi E, et al. Diagnostic criteria for small fibre neuropathy in clinical practice and research. Brain. 2019;142(12):3728-36.

5. Koch F, Kampgen E, Schuler G, Romani N. Isolation, enrichment, and culture of murine epidermal langerhans cells. Methods Mol Med. 2001;64:43-62.

6. Lauria G, Bakkers M, Schmitz C, Lombardi R, Penza P, Devigili G, et al. Intraepidermal nerve fiber density at the distal leg: a worldwide normative reference study. J Peripher Nerv Syst. 2010;15(3):202-7.
7. George DS, Jayaraj ND, Pacifico P, Ren D, Sriram N, Miller RE, et al. The Mas-related G protein-coupled receptor d (Mrgprd) mediates pain hypersensitivity in painful diabetic neuropathy. Pain. 2024.
8. Zhang H, Lecker I, Collymore C, Dokova A, Pham MC, Rosen SF, et al. Cage-lid hanging behavior as a translationally relevant measure of pain in mice. Pain. 2021;162(5):1416-25.
9. Hao Y, Stuart T, Kowalski MH, Choudhary S, Hoffman P, Hartman A, et al. Dictionary learning for integrative, multimodal and scalable single-cell analysis. Nat Biotechnol. 2024;42(2):293-304.
10. Jin S, Guerrero-Juarez CF, Zhang L, Chang I, Ramos R, Kuan CH, et al. Inference and analysis of cell-cell communication using CellChat. Nat Commun. 2021;12(1):1088.
